# Exploring the connections between digestion and detoxification in microevolution of insecticide resistance of the tea tortrix moth, *Adoxophyes honmai*

**DOI:** 10.1101/2025.06.18.660450

**Authors:** T. M. Lee, W. A. Nelson, C. D. Moyes

**Affiliations:** Department of Biology, Queen’s University, Kingston, ON, Canada, K7L 3N6

**Keywords:** phenotypic plasticity, microevolution, multi-drug resistance, lipase, cytochrome p450

## Abstract

While the evolution of insecticide resistance is often assumed to come with a fitness cost, there are instances where insect populations that acquired resistance fail to show an evolutionary cost to maintaining the trait in the absence of insecticide. In comparing two populations of tea tortrix moth, *Adoxophyes honmai*, we found the absence of a cost of resistance but also noted differences in digestive enzyme gene expression. This raised the possibility that insecticide resistance coevolved with enhanced digestive capabilities, potentially offsetting putative costs of resistance. This study explored gene transcript patterns that may influence how traits manifest spatially and temporally, evaluating potential connections between digestion and the costs of resistance. We found that our resistant larvae had constitutively greater transcript levels of multiple putative digestive genes as well as a marker of resistance, CYP9A170. The putative digestive genes were expressed mostly in the digestive tract, whereas the tissue-specific pattern of CYP9A170 expression was strain-dependent. For most genes, the difference in expression between susceptible and resistant larvae remained consistent throughout development. Interestingly, the expression of an ABC transporter was upregulated in response to tebufenozide exposure, but only in the resistant larvae. A comparison of *A. honmai* transcriptomes suggests that the majority of differentially expressed genes between populations may not be directly contributing to resistance, but rather microevolutionary variations specific to individual populations. Future studies on fitness costs of resistance should consider other physiological systems and their interactions with direct mechanisms of resistance.

## Introduction

Insecticide resistance evolves in a population when individuals that are less affected by insecticide have a selective advantage and higher fitness [1]. Most cases of insecticide resistance arise through two major mechanisms: target site insensitivity and enhanced toxin metabolism [2, 3]. In the case of target site insensitivity, a mutation in the target of an insecticide reduces binding of the insecticide to its target site. For example, mutations on acetylcholinesterase and voltage-gated sodium channels cause these proteins to be less perturbed by pesticides, specifically organophosphates and pyrethroids respectively [4, 5]. Alternatively, insects may evolve resistance through metabolic means with superior detoxification pathways or improved excretion of insecticides from cells and organisms. Key metabolic enzymes that are often implicated and upregulated in the evolution of resistance include cytochrome p450s (CYPs) [6–8], glutathione-S-transferases (GST) [9], esterases, and ATP-binding cassette (ABC) transporters such as multi-drug resistant (MDR) transporters [10, 11].

Evolved differences in resistance may manifest along different timelines and life stages, with or without exposure to insecticide. For example, resistant tea tortrix populations in the field differed in terms of the transcription of genes associated with resistance, namely CYPs and esterases [12]. Since these transcriptional differences persisted in the absence of treatment, it is likely that mutations affected basal expression. However, there is also the potential for resistance arising from differences in inducibility of defensive genes, which become evident only in response to treatment [13].

The evolution of resistance through these mechanisms is often assumed to come with fitness costs. For example, in *Drosophila melanogaster* a mutation in their GABA receptor confers resistance to cyclodiene but causes paralysis under high temperatures [14]. In other cases, the mechanisms that confer resistance may increase energetic costs. For instance, mosquitoes *Culex pipiens* that overexpressed esterases had less energy storage than susceptible mosquitoes [15]. In most cases, it is unclear how the resistance mutation disrupts energy balance, and it is frequently assumed that resistance comes with a metabolic cost that might put the individual at a deficit. However, whether fitness costs are present depends on a plethora of factors, including the resistance mechanism, the strain in question, its evolutionary history, and their interactions [16, 17].

The smaller tea tortrix, *Adoxophyes honmai*, is a major pest of tea in Japan. These moths have developed resistance to multiple insecticides [18, 19]. Uchibori et al. [12] showed that tebufenozide resistance was associated with enhanced metabolism and target site insensitivity arising from a mutation on the ecdysone receptor. In a recent study, we sought to assess the fitness costs of resistance in these moths and surprisingly found that the evolution of tebufenozide resistance did not appear to come with fitness costs [20]. The level of tebufenozide resistance, measured in LD_50_, among *A. honmai* families did not have a negative correlation with their fitness-proxy life-history traits, development time and pupal weight [20]. Where the costs of resistance are largely energetic, there is a potential of overcoming such costs through evolutionary adaptations that improve digestion and nutrient assimilation. Indeed, resistant larvae developed slightly slower and produced heavier pupae when food was abundant [20]. Survival rate of resistant pupae was also markedly higher than susceptible pupae when food was limited [20], which suggested that the resistant population possessed other traits that would manifest in fitness advantage over the susceptible population under certain conditions.

Although we found no evidence of a fitness cost of tebufenozide resistance [20], we noted in a preliminary study many differences between susceptible and resistant lines in genes associated with digestion and intermediary metabolism. These differences could, in priniciple, offset energetic deficits arising from resistance, explaining why we found no costs of resistance between these strains.

In this study, we further explored the differences in detoxification and digestive enzyme genes between susceptible and resistance *A. honmai* larvae. Our focus was on differences that emerge at the transcript level, recognizing that such differences may arise from mutations acting on genes in cis or trans. We focused on capturing the transcript variation seen between these lines (i) at different life stages, (ii) under different dietary conditions, and (iii) in response to insecticide treatment. For example, the magnitude of transcript differences between lines may depend upon larval stage; if differences are diminished at a particular larval stage, this could correspond to a stage-specific vulnerability in the resistant population. With respect to dietary conditions and insecticide exposure, transcript profiles show if resistant animals have greater basal expression, and/or whether they are able to induce expression more rapidly or to a greater extent than susceptible populations. The time course of gene expression is also an important experimental design consideration because some differences between lines may only emerge upon extended exposure. Though transcript-based studies cannot demonstrate differences in protein activity or function in the context of physiological systems, they can reveal genetic differences between susceptible and resistant lines that may contribute directly to resistance or indirectly to overcome the costs of resistance. In this regard, we use the term “expression” to refer specifically to mRNA patterns.

## Methods

### Animals

The stock animals, two strains of *Adoxophyes honmai*, were maintained as described in Lee et al. [20]. The susceptible (S) was imported from the National Institute for Agro-Environmental Sciences Japan in 2013. It had been contained in Japanese laboratories since the 1960s, and is considered naïve to insecticides. The resistant strain (R) was collected and imported in Spring 2019 from Kanaya, Japan. It is known to be resistant to tebufenozide [20]. All animals were fed an artificial silkworm diet (SilkMate 2M; Nosan Corporation, Yokohama, Japan) *ad libitum* under a light-dark cycle of 14:10 h. Animals in the colony were raised communally, while experimental animals were isolated within 24 h of hatching into 1.5 oz (43 mL) translucent plastic cups (Dart Container P150N).

### RNA-seq on midgut tissue

Fifth instar larvae were dissected for their midgut to perform RNA-seq. Larvae of either strain were isolated from food during their molt to fifth instar and were given food *ad libitum* within 24 hours of successful molting. Larvae were then dissected three days after food access to collect their midgut. The midguts from 4 larvae of the same sex were pooled into each biological replicate. Samples were stored in RNALater solution at -80°C until further analysis.

To construct libraries for RNA-seq, total RNA was extracted from samples with TRIzol Reagent (Thermo Fisher) following manufacturer’s instruction. The integrity of extracted RNA was assessed using gel electrophoresis and Agilent Bioanalyzer. The total RNA of 5 susceptible and 3 resistant biological replicates were then treated with DNase (DNA-free^TM^, Thermo Fisher AM1906) before strand-specific library construction (NEBNext® Ultra^TM^ II Directional RNA Library Prep kit for Illumina®, NEB E7760; NEBNext® Multiplex Oligos for Illumina®, NEB E6440S).

Sequencing was performed by The Centre for Applied Genomics, The Hospital for Sick Children, Toronto, Canada. The libraries were sequenced on an Illumina NovaSeq 6000 SP flowcell using the paired-end 100-bp mode. Sequencing data were then processed using the standard Illumina pipeline and Fastq files were generated for data processing and assembly.

### Tissue-specific pattern in gene expression

Larvae of either strain (8 susceptible; 9 resistant) were dissected one day after molting into fifth instar to explore the expression of genes of interest in different tissues. The entire digestive tract (DT) was separated from the rest of the carcass. Sex of the larvae was also noted. Specimen was flash frozen and stored at -80°C until further analysis. The proportion of gene expression detected in the DT was calculated based on the concentrations of RNA extracted from either DT or the carcass using this formula: (Expression_DT_ * [RNA_DT_]) / (Expression_DT_ * [RNA_DT_] + Expression_Carcass_ * [RNA_Carcass_]). In general, the digestive tract accounted for about 20 % of the mass of the entire animal, and about 50 % of the RNA of an animal came from its digestive tract.

### Inducible gene expression in response to food intake

To test whether the two strains of larvae alter gene expression after immediate feeding, third and fifth instar susceptible and resistant larvae were allocated into “unfed” or “fed” conditions. All larvae were isolated from food during their molt into either third or fifth instar. Within 24 hours of isolation from food, larvae were weighed and given either no food for control or food *ad libitum* for the fed condition. In total, there were 9 susceptible unfed, 10 susceptible fed, 8 resistant unfed, and 8 resistant fed 3^rd^ instar larvae. There were 9 5^th^ instar larvae in each of the four strain-feeding conditions. All larvae were weighed then flash frozen 4 hours after the start of feeding and stored at -80°C until further analysis.

### Developmental pattern in constitutive gene expression

To investigate how gene expression patterns change differently between strains throughout development, whole-body samples of susceptible and resistant *A. honmai* were collected during their five stages of larval development. Animals were checked every day for signs of molting and sampled within 24 h of molting into a new larval stage. Animals were also sampled at the midpoint of each of the five larval stages, at 2, 2, 1, 1, and 3 days after molting respectively, for a total of 10 developmental time points. Because the early larvae had smaller body size, each biological replicate of first and second instar larvae consisted of a pool of 10 and 3 individuals respectively. Replicates of third to fifth instar larvae consisted of one individual each. There were 6 biological replicates in each of the 20 strain-development stage combinations. All samples were flash frozen and stored at -80°C until further analysis.

### Inducible gene expression in response to tebufenozide

To test whether the two strains respond differently to tebufenozide exposure in gene expression, third instar susceptible and resistant larvae were given droplets that contain different amounts of tebufenozide. All larvae were isolated from food during their molt into third instar. Within 24 hours of isolation from food, larvae of either strain were given a 0.4 µL droplet of water [21, 22], which contained 0 ng, a low dose of 1.2 ng, or a high dose of 8 ng of tebufenozide (Confirm 2F, Gowan, Yuma, AZ, USA; Lot# CJBAL7001). The low and high doses induced approximately 36 % and 82 % lifetime mortality respectively in exposed susceptible larvae; however, the lowest dose tested that induced mortality in the resistant larvae was greater than 8 ng (60 ng; Figure S1). Exposed larvae were monitored to ensure that the droplet was fully ingested within 2 hours. Only larvae that completed ingested their droplets were included in the experiment. Larvae were sampled 2, 4, and 6 hours after exposure, flash frozen, and stored at - 80°C until further analysis. There were 3 biological replicates for each of the strain-dose-time combinations.

### qPCR

Total RNA was extracted using the RNeasy Plus Micro Kit (QIAGEN Cat# 74034) following the manufacturer’s instruction for gene expression experiments. The integrity and quality of the extracted RNA were inspected using gel electrophoresis and 260/280 nm absorbance respectively. One microgram of RNA was then reverse transcribed into cDNA using LunaScript® RT SuperMix Kit (NEB E3010), and diluted to 5 ng/µL for storage at -20°C.

For qPCR, the amplification efficiencies of primers (Table S1) were verified using thermal gradients and standard curves. Genes of interest related to digestion and detoxification, including amylase, trypsin, chymotrypsin, lipase, and a multi-drug resistance 49-like protein (mdr49), were identified based on the RNA-seq experiment, the sequences from which were used to design primers. Primers for *CYP9A170*, another gene of interest associated with tebufenozide resistance, and *rp49*, the reference gene, was obtained from Uchibori-Asano et al. [12]. The stability of rp49 was validated against actin and ecdysone receptor in a preliminary experiment. The expression of hormone receptor 3 (HR3) was measured as an indicator for responsiveness to tebufenozide [23]. For qPCR, reaction mixes consist of 5 µL of the diluted cDNA, 0.5 µL each of 10 µM forward and reverse primers, 4.5 µL of nuclease-free water, and 10 µL of Luna® Universal qPCR Master Mix (NEB M3003). Reactions were performed in CFX96 (Bio-Rad) with the following conditions: 60 s at 95°C, 40 cycles of 15 s at 95°C, 30 s at 58 °C and plate read, followed by a melt curve from that increases in 0.5°C increments every 5 s from 65°C to 95°C. Each biological replicate was assayed with technical duplicates. No signal was detected in no template or no reverse transcriptase controls. Expression of genes of interest was normalized using the expression of *rp49*.

### Transcriptomic Analysis of Published RNA-seq data

RNA-seq datasets previously published by Uchibori-Asano et al. [12] were downloaded from NCBI (DRX132508, DRX132509, DRX132510, DRX132511, DRX132512, DRX132513, DRX132517, DRX132518, DRX132519, DRX133886, DRX133887, DRX133888). These

RNA-seq data came from *A. honmai* with different origin and levels of resistance to tebufenozide. Transcripts in these transcriptomes were quantified using the transcriptome assembly constructed with our RNA-seq data (see Data Analysis). After that, the expression of genes that were implicated in the differential gene expression analysis of our RNA-seq data and other experiments were examined.

### Data Analysis

#### RNA-seq

RNA-seq reads were inspected for quality control and preprocessed using HTStream (https://github.com/s4hts/HTStream; accessed September 23, 2021), including removal of adapter sequences, contaminants, PCR duplicates, poly A/T tails, and reads shorter than 50 bp, as well as filtering out low quality reads and extracting longest subsequence with no Ns. The reads were then used to assemble a transcriptome *de novo* using Trinity 2.14.0 [24]. The assembly was annotated using Trinotate 4.0.0 [25]. After that, transcripts in each library were quantified at the gene and transcript levels using the assembly and salmon 1.10.2 [26] in Trinity. Normalized values were expressed as TMM (trimmed mean of M values)-normalized transcript per million (TPM). Pearson correlation between all replicates was calculated. The raw counts at the gene level were then used for differential expression analysis with *limma* (voom) [27]. Finally, differentially expressed genes were filtered with the criteria of a minimum 4-fold difference and *p* < 0.001, followed by a gene ontology (GO) enrichment analysis on the filtered genes.

#### Statistics

To determine the effects of different variables (strain, tissue, developmental stage, food intake, time, and tebufenozide treatment) on gene expression, we compared linear models (*lm()* in R) [28] that accounted for the main and interaction effects of these independent variables when applicable. A null model that contained no independent variable was also included for comparison. We used the second order Akaike’s information criterion (AICc) to identify the best-fit model for each dataset [29]. If the best-fit model included a main effect of an independent variable that had multiple levels or an interaction between multiple variables, *post hoc* pairwise comparisons were conducted based on the best-fit model using Tukey tests with Benjamini-Hochberg adjustment for multiple testing to identify significant differences.

## Results

### Differential gene expression between strains in the midgut

Figure 1A summarizes the expression patterns of differentially expressed genes (DEGs) in the midguts of 5^th^ instar susceptible and resistant larvae. Because biological replicates differed more by strain than sex (Fig S3), we pooled replicates of different sexes for downstream differential gene expression analysis by strains. Biological replicates were similar within each strain with high correlation (average Pearson correlation between replicates: 0.90; lowest Pearson correlation: 0.84). Out of 62,025 genes in the assembly comparing the midguts of susceptible and resistant larvae, we identified 1,855 DEGs that passed the filter criteria for 4-fold difference and false discovery rate of 0.001 (Fig 1A). Amongst these, 1,746 DEGs were upregulated, and 109 were downregulated in the resistant larvae compared to the susceptible larvae.

**Figure 1.**
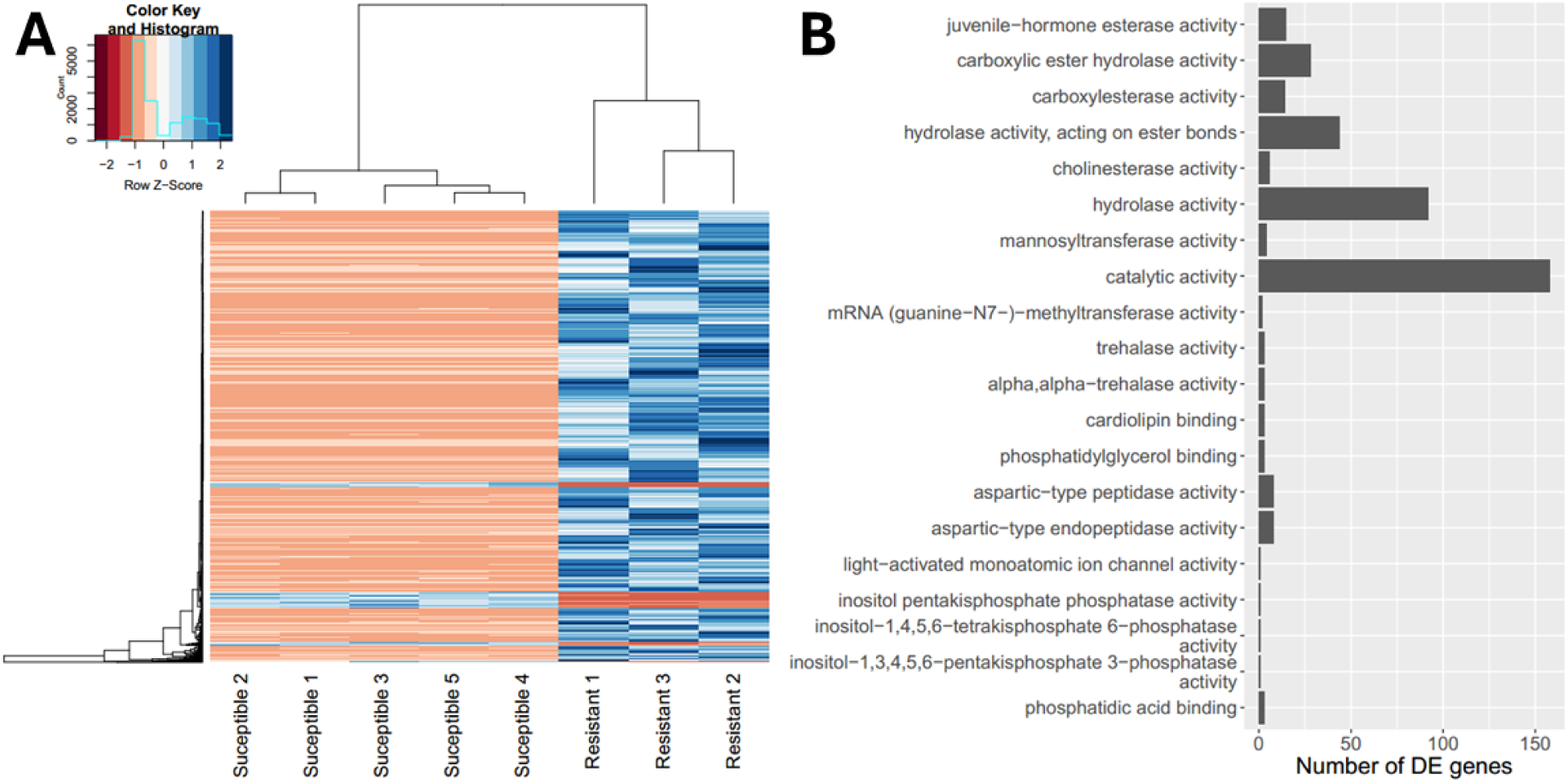
Differential gene expression analysis of midguts in susceptible and resistant larvae. Each biological replicate of midguts consisted of a pool of 4 fifth-instar larvae that had been fed *ad lib* for three days. RNA was analyzed by RNA-seq. (A) Heatmap of differentially expressed (DE) genes. (B) Top 20 enriched “biological process” gene ontology categories with the number of DE genes in each category.

The GO enrichment analysis identified 345 categories that were enriched in the DEGs. Amongst these categories, 198 categories had the terms “biological process”, 93 had “molecular function”, and 34 had “cellular process”. Sorted by their p-value for overrepresentation, the top 20 GO categories for biological process contain those related to detoxification and digestion, such as “carboxylesterase activity” and “trehalase activity” (Fig 1B).

From this analysis, we noted a number of genes of obvious relevance (Supplementary Information B). Resistant insects had elevated expression of CYPs and glutathione S-transferase, consistent with modes of metabolic insecticide resistance. The resistant larvae also had elevated levels of key digestive enzymes, including trehalase, peptidases and a homolog of vertebrate pancreatic lipase. Additionally, resistant larvae had greater expression of various esterases, which may be involved in either detoxification or digestion.

Further analysis focused on CYP9A170 as an indicator of resistance because the difference in its expression between susceptible and resistant larvae was the greatest among the CYPs associated with tebufenozide resistance in *A. honmai* [12], which we tested in a preliminary experiment (Fig S2). Because DEGs potentially related to digestion were upregulated in resistant larvae, we also performed a literature search for confirmed digestive enzymes for carbohydrates, peptides, and lipids in insects [30–33]. Taking into consideration the literature search, the DEG analysis, and fold differences in expression between susceptible and resistant midgut of all the annotated genes in our transcriptome, we identified amylase, maltase, trypsin, chymotrypsin, gastric lipase, and pancreatic lipase as genes of interest associated with digestion.

### Tissue-specific pattern in gene expression

Some of the genes we identified for analysis were annotated based on their vertebrate homologs. Thus, it seemed prudent to verify that the expression of putative digestive enzyme genes was localized to the digestive tract. Also, insect RNA analysis is often complicated by their small size, leading to analyses of the whole animal, which becomes an issue when the tissue of significance is a smaller fraction of the total mass. This raises a concern about real differences in a tissue lost when pooling all the RNA in a whole-body sample. By independently analyzing the digestive tract and the carcass, we also reduced the potential influence of a largely negative signal in the carcass diminishing the signal derived from the smaller (by mass) digestive tract.

Figure 2A summarizes the tissue distribution of our genes of interest reported from both susceptible and resistant fifth-instar larvae. As expected, almost all (mean > 96 %) of qPCR-based signal for each of the digestive genes were detected in the digestive tract. However, from the perspective of the whole body, only about 40 % of CYP9A170 expression was found in the digestive tract (Fig 2A).

**Figure 2.**
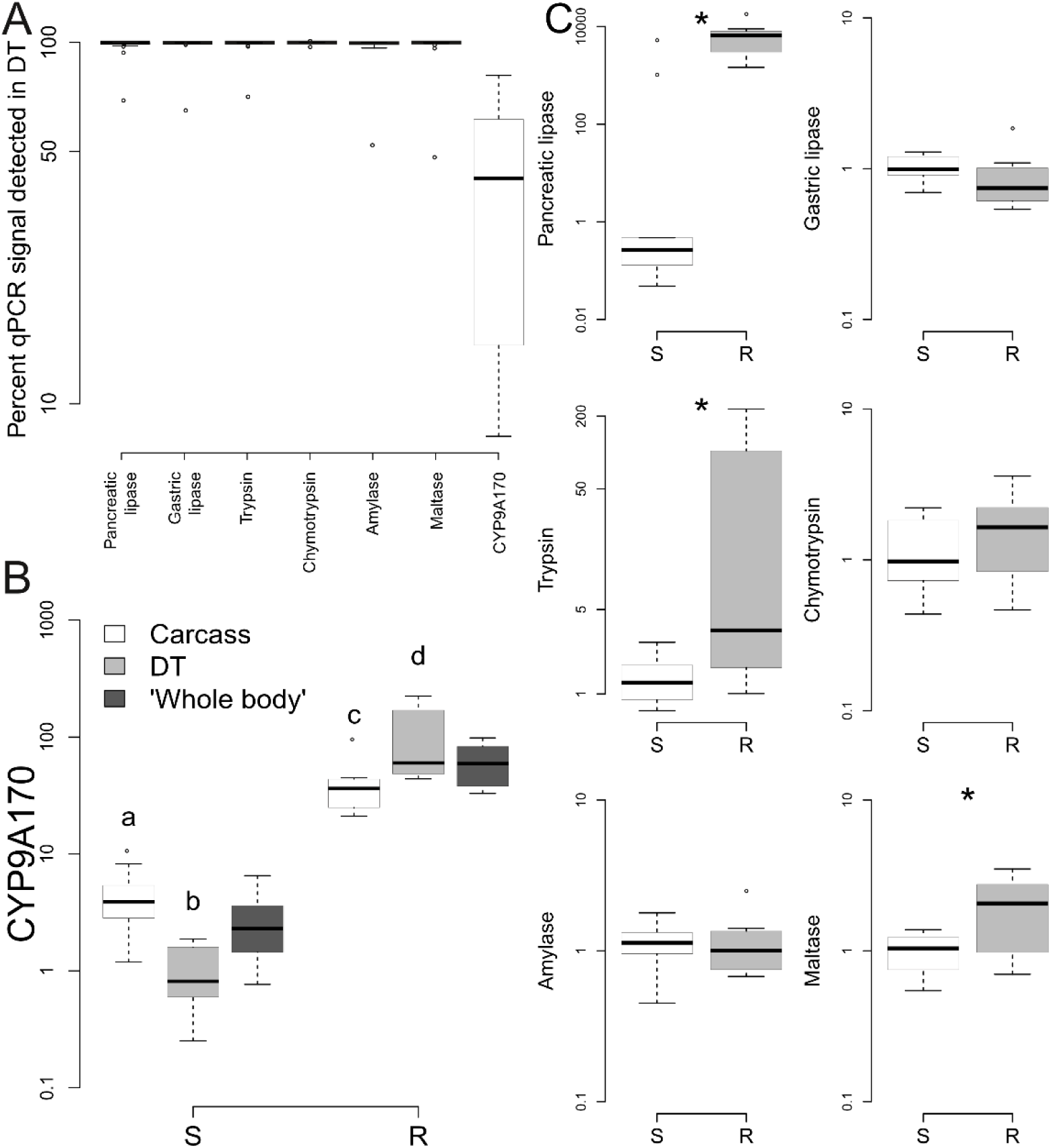
Tissue-specific gene expression patterns for putative digestive enzymes and CYP9A170 in the digestive tract (DT) and the rest of the carcass of fifth instar susceptible (S) and resistant (R) larvae. For each sample, Cq values of each gene were first normalized using the Cq of the reference gene (rp49), then divided by the mean normalized value of the DT of susceptible larvae to obtain the normalized relative fold-change for each gene. Susceptible: n = 8; resistant: n = 9. (A) Percent of qPCR-based transcript levels detected in the DT for both S and R individuals based on the RNA concentration extracted from both the DT and the rest of the carcass. Each value represents one individual and was calculated using this formula: (Expression_DT_ * [RNA_DT_]) / (Expression_DT_ * [RNA_DT_] + Expression_Carcass_ * [RNA_Carcass_]). (B) Normalized fold-change in CYP9A170 expression in the DT, the rest of the carcass, and the expected value in a whole-body specimen calculated using the DT and carcass values, relative to the mean in susceptible DT. Different letters indicate significant differences (p < 0.05) identified in the *post hoc* pairwise comparisons using Tukey’s test with Benjamini-Hochberg adjustment for multiple testing. (C) Normalized fold-change in gene expression of putative digestive enzymes in the DT, relative to the means in susceptible DT. Asterisks indicate significant differences between strains in the DT based on model selection with AICc.

Figure 2B reports the tissue distribution for CYP9A170 expression. Interestingly, the model of best fit identified an interaction between strain and tissue on the expression of CYP9A170 (Table S2). In both carcass and digestive tract, the mRNA levels of CYP9A170 were much higher in the resistant larvae than susceptible larvae: 10-fold greater in the carcass, 100-fold greater in the digestive tract. Within strains, the relative amounts of CYP9A170 also differed (Table S3). Susceptible insects had 4-fold greater expression in the carcass than the digestive tract. The pattern was reversed in resistant animals, with about 2-fold more mRNA in the digestive tract. This strain-specific differential expression between tissues would not be detected had this analysis been performed on whole animals. By subdividing the animals, we revealed that the mRNA for CYP9A170 was relatively enriched in the digestive tract of resistant larvae.

Figure 2C summarizes the digestive enzyme mRNA data exclusively from the digestive tract. The mRNA levels of digestive genes were much lower in the carcass, frequently bordering the levels of detection and therefore less reliable (Table S4). We found that the expression of pancreatic lipase, trypsin, and maltase were significantly different between the susceptible and resistant strains (Table S5). In comparison to susceptible larvae, resistant larvae expressed 10-fold more pancreatic lipase, 38-fold more trypsin, and 2-fold more maltase in the digestive tract (Fig 2C). Only trypsin expression differed between sexes (Table S5), where resistant females expressed more trypsin (Fig S4).

The fold differences seen in this study did not fully agree with our transcriptome data in several aspects. While they were analyzed on different insects, we also note that potential differences could have arisen from variations in both developmental stage (early vs mid 5^th^ instar) and time since feeding. The next studies addressed the potential influence of developmental stage and feeding on gene expression.

### Food intake induced gene expression

Many digestive enzymes are induced by feeding, so our next study focused on how time-since-feeding affected expression of our genes of interest. We conducted these analyses at two stages: 3^rd^ and 5^th^ instar, using whole animals. Based on Figure 2A, it is clear that, for these genes, most of the mRNA in a whole animal arises from the digestive tract itself. At these stages, the animals are large enough to provide enough mRNA to analyze individuals.

Figure 3 shows how feeding affected digestive enzyme expression, and whether the pattern differed between strains. In general, there were no increases in mRNA for any enzyme as a result of feeding. Pancreatic lipase, which was about 1000-fold different between strains when measured in the digestive tract a day after molting (Fig 2C), showed little difference between strains when measured within one day of molting into 3^rd^ instar without feeding (Fig 3A), and a 4-fold difference between strains after molting into 5^th^ instar (Fig 3D). Interestingly, 3^rd^ instar resistant larvae expressed about 4-fold more pancreatic lipase only after 4 h of feeding (Fig 3A). The converse was seen with trypsin. Although the data were variable, the strain difference in trypsin in the digestive tract of 5^th^ instar larvae was about 38-fold (Fig 2C), whereas differences between strains in this study were as much as 50-fold in either 3^rd^ or 5^th^ instars (Fig 3B, 3E). Resistant larvae also had about 2-fold greater expression of amylase than susceptible larvae in 5th (Fig 3F) but not 3^rd^ instar (Fig 3C).

**Figure 3.**
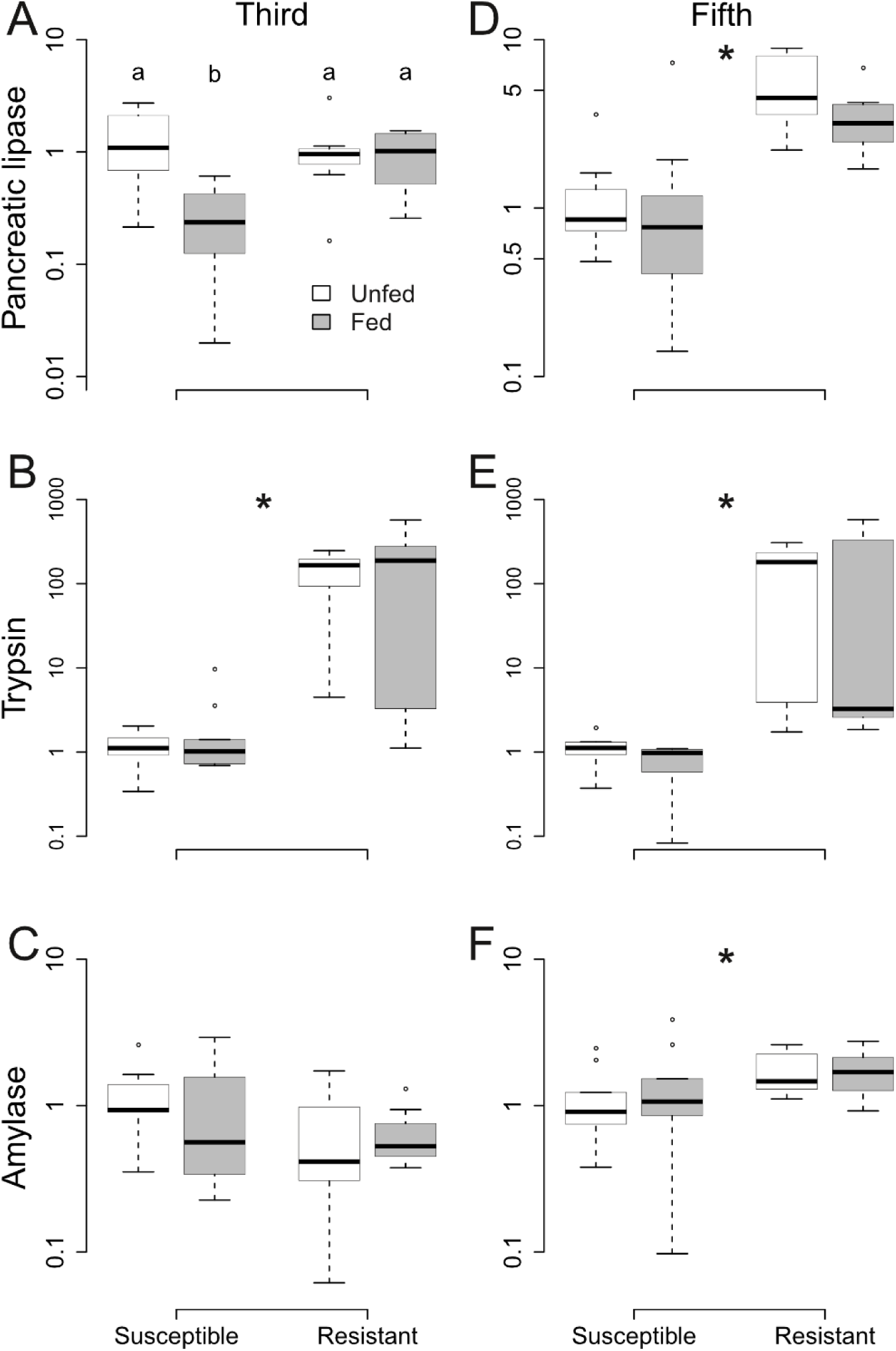
Normalized relative fold-change in gene expression of putative digestive enzymes in whole-body samples of third and fifth instar susceptible and resistant larvae that were either fed or unfed for 4 hours. Expression values of each gene for each sample were normalized using the Cq of the reference gene, rp49, then divided by the mean normalized value of starved susceptible larvae for the respective larval instar of the respective gene to obtain the normalized relative fold-change. Third instar: susceptible unfed n = 9, susceptible fed n = 10, resistant n = 8; fifth instar: n = 9. Asterisks indicate a main effect of strain on gene expression identified by the model of best-fit with AICc. Different letters indicate significant differences (p < 0.05) identified in the *post hoc* pairwise comparisons using Tukey’s test with Benjamini-Hochberg adjustment for multiple testing.

The supplementary information summarizes the analysis of the entire dataset, reporting strain effects, feeding effects and interactions (Tables S6, S7). Collectively, the studies summarized in Figures 2 and 3 emphasize the influence of instar and time-since-moult in determining the magnitude of differences in digestive enzymes.

### Developmental patterns for constitutive gene expression

Given the potential influence of developmental stage on gene expression, we investigated the expression profiles of the genes of interest throughout larval development (Fig 4). Resistant larvae had greater constitutive expression of all measured genes except amylase (Fig 4B) than susceptible larvae. The fold difference in expression between strains varied for different genes though. Compared to susceptible larvae, resistant larvae expressed approximately 7-fold more pancreatic lipase (Fig 4A), 53-fold more trypsin (Fig 4C), 3-fold more maltase (Fig 4D), 26-fold more CYP9A170 (Fig 4E), and 2-fold more mdr49-like protein (Fig 4F). However, the magnitude of between-strain differences in gene expression did not vary with developmental stage, as there was no interaction between strain and developmental stage (Table S8).

**Figure 4.**
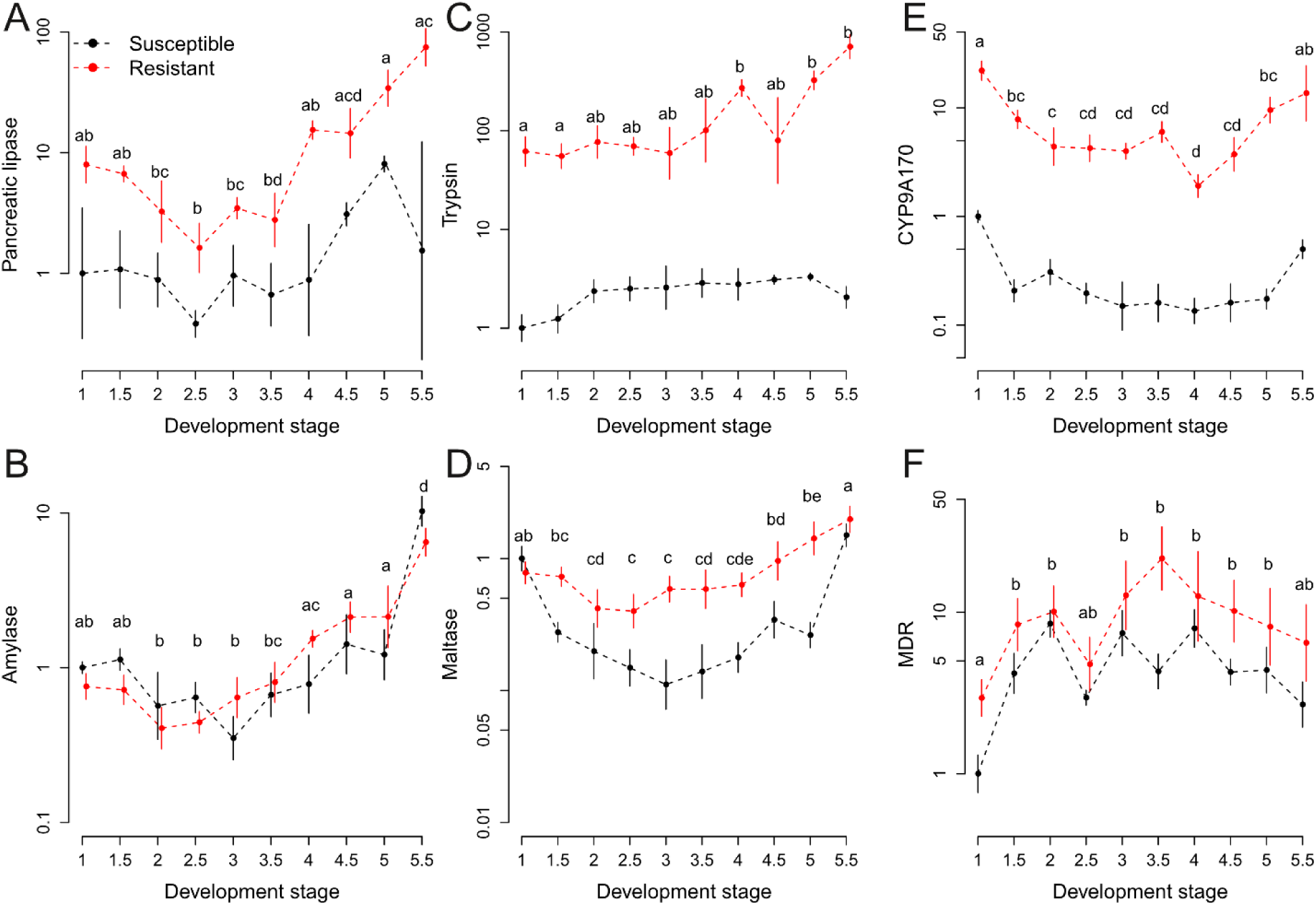
Normalized relative fold-change in gene expression (mean ± SE) of (A – D) putative digestive enzymes and (E – F) detoxification-related proteins in susceptible and resistant larvae of different development stages. Expression values of each gene for each sample were normalized using the Cq of the reference gene, rp49, then divided by the mean normalized value of first instar susceptible larvae of the respective gene to obtain the normalized relative fold-change. n = 6 in each strain-stage combination. Different letters indicate significant differences between development stages in the *post hoc* pairwise comparisons using Tukey’s test with Benjamini-Hochberg adjustment for multiple testing.

The expression of digestive genes increased steadily through larval development before reaching a peak in the fifth instar (Fig 4A – D), although the expression of pancreatic lipase (Fig 4A), amylase (Fig 4B), and maltase (Fig 4D) did experience a dip around the second and third instars. In contrast, the expression of detoxification related genes appeared to be more consistent throughout larval development (Fig 4E – F). The expression of CYP9A170 was greater in the first and fifth instar (Fig 4E), while the expression of MDR was lower only in the first instar hatchlings (Fig 4F). The comparisons between developmental stages for each investigated gene were summarized in Tables S9 – 14.

### Tebufenozide exposure induced gene expression

Because we explored the influence of feeding on digestive gene expression, it seemed prudent to also examine the effect of insecticide exposure on the expression of detoxification genes. Figure 5 reported both constitutive and tebufenozide-induced expression of a positive control for tebufenozide exposure, HR3, as well as detoxification genes, namely CYP9A170 and mdr49-like protein. The expression of these three genes demonstrated distinctive patterns in response to tebufenozide in relation to either strain or dosage.

**Figure 5.**
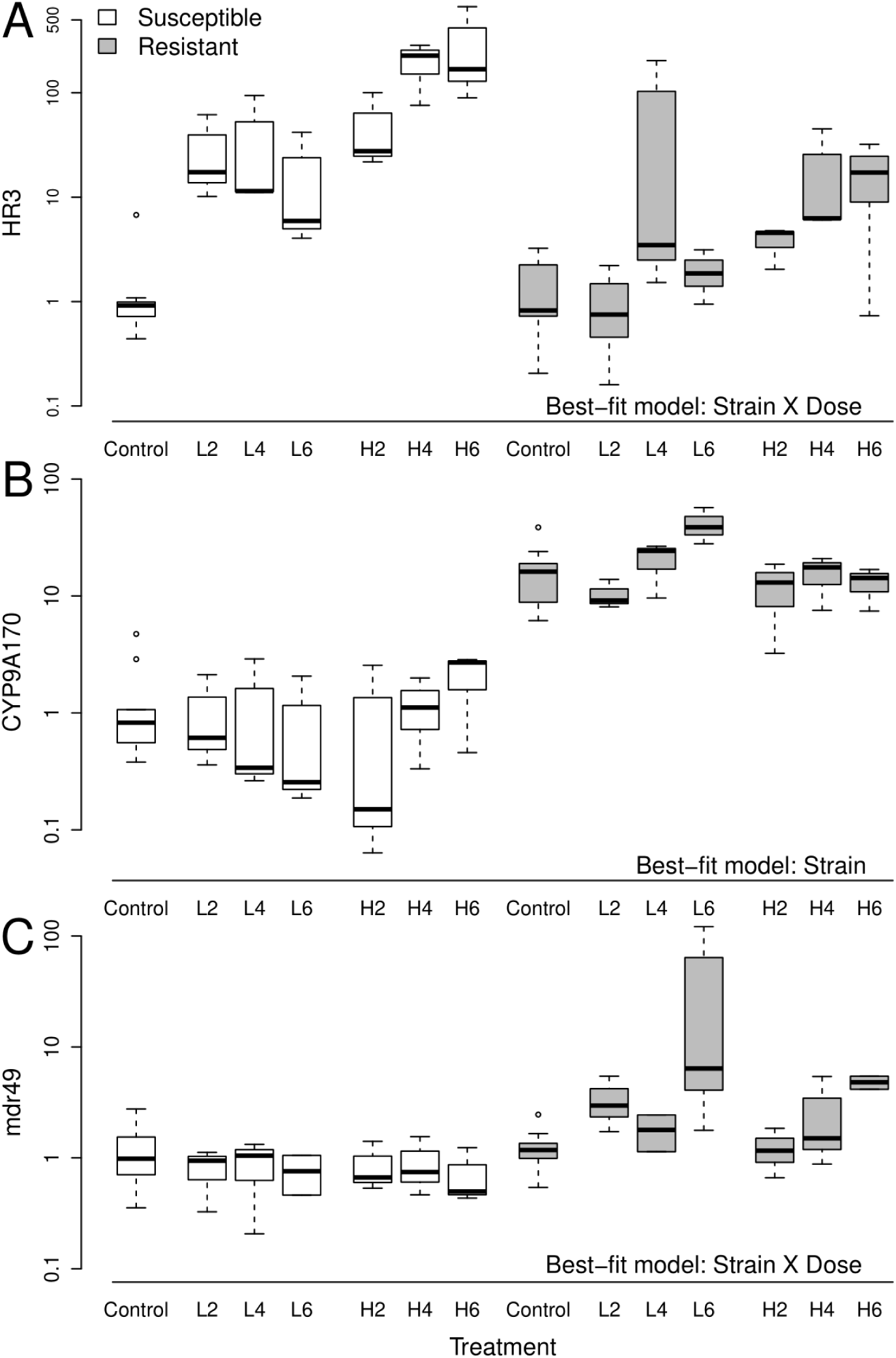
Normalized relative fold-change in gene expression of (A) HR3, (B) CYP9A170, and (C) mdr49-like protein in susceptible and resistant third instar larvae that were exposed to different doses (control: 0 ng; low (L): 1.2 ng; high (H): 8 ng) of tebufenozide for 2, 4, or 6 hours. n = 3 in each group. Expression values of each gene for each sample were normalized using the Cq of the reference gene, rp49, then divided by the mean normalized value of the respective gene for control susceptible larvae given water to obtain the normalized relative fold-change. Data for control larvae exposed to water for each of the 3 durations were not significantly different and were pooled for presentation. The best-fit model for each data set chosen based on AICc is indicated for each gene.

Susceptible and resistant larvae did not differ in their constitutive expression of HR3, but both strains responded in a dose-dependent manner to tebufenozide, albeit to different magnitudes (Fig 5A). Larvae generally expressed more HR3 when they were exposed to a higher dose of tebufenozide compared to a low dose. However, susceptible larvae had a greater increase in HR3 expression than resistant larvae at either tebufenozide dosage, reflected in the interaction term between strain and dose in the model of best fit (Table S15). HR3 expression was comparable between susceptible larvae exposed to 8 ng tebufenozide and resistant larvae exposed to 1.2 ng tebufenozide.

In contrast, resistant larvae had greater expression of CYP9A170 than susceptible larvae regardless of tebufenozide exposure (Fig 5B). Resistant larvae had about 20-fold greater expression of CYP9A170 than susceptible larvae when they were either given water or exposed to one of the two doses of tebufenozide. This between-strain difference in CYP9A170 expression is comparable to previous experiment (Fig 4E).

While the two strains had similar mdr49 expression in untreated animals, only resistant larvae increased expression of mdr49-like protein in response to tebufenozide (Fig 5C). Susceptible larvae showed no induction of this gene at any dose or time point. Contrarily, resistant larvae increased mdr49 expression by 2- to 4-fold after tebufenozide exposure. Because only resistant and not susceptible larvae expressed more mdr49 after tebufenozide exposure, the interaction term between strain and tebufenozide dosage was included in the model of best fit for mdr49 expression (Table S15). While mdr49 expression in resistant larvae appeared to increase to about 5- to 13-fold by 6 hours after tebufenozide exposure, time was not a factor included in the best-fit model (Table S15). The supplementary information also contains a detailed summary for *post-hoc* comparisons regarding the interaction between strain and dosage for both HR3 and mdr49 (Table S16 – S17).

### Expression profile in other A. *honmai* transcriptomes

The distinction between susceptible and resistant strains in the previously published RNA-seq libraries was not as clearcut. Based on the multidimensional scaling analysis of overall expression, the two resistant strains, Met and Yui, were distant from each other (Fig 6A). The Yui replicates were closer instead to replicates of the two susceptible strains, Kanaya and Haruno.

**Figure 6.**
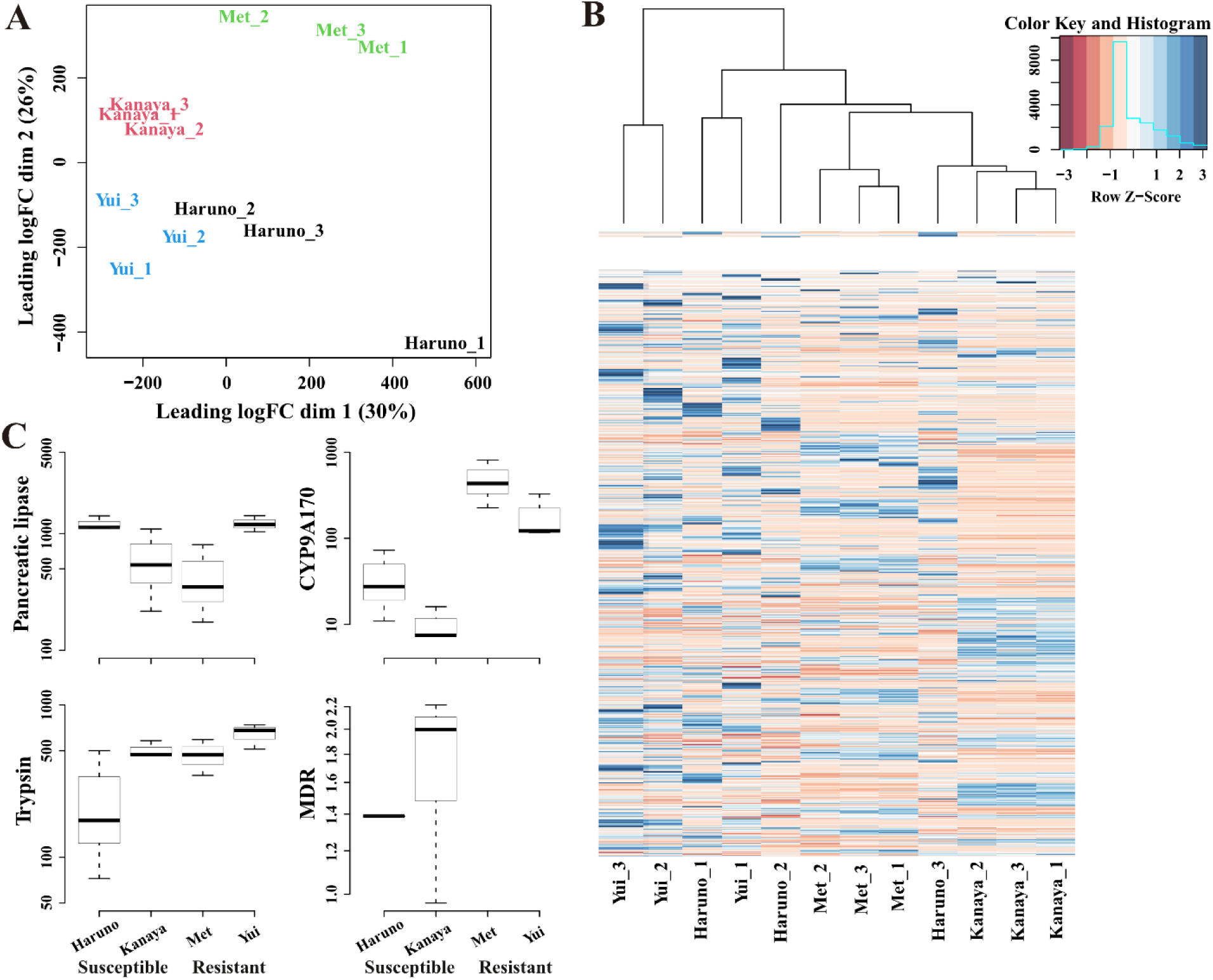
Analysis of previously published susceptible (Haruno and Kanaya) and resistant (Met and Yui) *Adoxophyes honmai* transcriptomes (Uchibori-Asano et al. 2019). (A) Multidimentional scaling (MDS) plot visualization of the distance between the transcriptomes of the 4 different strains of *A. honmai*. (B) Heatmap of the expression of the differentially expressed genes identified in our midgut transcriptomes (Fig 1). (C) TMM (trimmed mean of M values)- normalized transcript per million (TPM) of pancreatic lipase, trypsin, CYP9A170, and mdr49-like protein.

Focusing on the same 1,855 DEGs that were identified in our RNA-seq data (Fig 1A), the four strains appeared to have distinct expression profiles (Fig 6B). The resistant Yui replicates appeared to be more enriched, having greater expression of several clusters of genes. Two replicates of the susceptible Haruno replicates also had similar expression pattern compared to the Yui replicates. In contrast, the Met and Kanaya strains had slightly muted expression profiles, with each strain expressing a different cluster of genes.

The expression of specific genes in these four strains reflected the broader scale patterns. While expression of pancreatic lipase appeared to higher in the resistant Yui strain, its expression was comparable to that of the susceptible Haruno strain (Fig 6C). Similarly, both Yui and Met had high expression of trypsin, but so did the susceptible Kanaya strain (Fig 6C). On the other hand, the two resistant strains of Yui and Met had greater expression of CYP9A170 compared to the susceptible Haruno and Kanaya strains (Fig 6C). Interestingly, the mdr49-like gene was not detected in the two resistant strains.

## Discussion

Insecticide resistance often evolves through a suite of mechanisms that includes target-site insensitivity and enhanced toxin metabolism, as is the case for *Adoxophyes honmai* [2, 3, 12]. While evolution of resistance is generally expected to come with life-history costs [16, 17], the fitness of a field-collected tebufenozide-resistant strain of *A. honmai* was not lower than a susceptible strain under various conditions [20]. In this study, we further characterized our two strains of *A. honmai* at the transcript level in relation to resistance but also to digestion, which may compensate for energetic requirements of improved toxin metabolism. We confirmed that our resistant strain had greater constitutive gene expression of detoxification enzymes associated with tebufenozide resistance and found that they upregulated expression of an ABC transporter in response to tebufenozide exposure. In addition, the resistant strain had greater expression of putative digestive enzymes throughout development. Finally, comparisons with previously published transcriptome of other *A. honmai* strains [12] suggest that different strains that are resistant to the same insecticide likely have microevolutionary variations other than the mechanisms that contribute directly to resistance. Such differences may interact with mechanisms of resistance to determine whether evolution of insecticide resistance comes with a fitness cost.

Gene ontology analysis of our midgut transcriptomes revealed that the resistant larvae were enriched in putative digestive genes, as well as detoxification genes typical of insecticide resistance. This observation hints at the physiological differences between our two *A. honmai* strains underlying their life-history differences. Lee et al. [20] previously reported that not only did resistant larvae develop slower and produce heavier pupae when food was abundant, they also had much greater survival when food was limited. To make connections between life-history and physiology, we decided to further characterize a selection of genes related to digestion and detoxification regarding tissue specificity, development, and responses to stimuli.

Putative digestive enzymes were expressed predominantly in the digestive tract. Much of digestion in insects occurs in their midgut [31]. Indeed, digestive amylase [34], protease [35, 36], and lipase [33] have been found to be expressed in the midgut of Lepidopterans. Our findings that the tested genes were similarly localized to the digestive tract therefore provide confidence to their involvement in digestion. On the other hand, CYP9A170 expression had a strain-specific tissue localization pattern (Fig 2); CYP9A170 was expressed more in other parts of the body in susceptible larvae, but more in the digestive tract in resistant *A. honmai*. CYPs involved in xenobiotic metabolism are usually expressed in tissues engaged in the first line of defense, which include the midgut, Malpighian tubules, and the fat body [8, 37]. Tissue-specific expression profile also depends on the specific CYP. For instance, *Spodoptera exigua* had at least five clusters of CYPs that had distinct expression profiles among different tissues [13]. Our findings suggest that CYP9A170 expression in the digestive tract may have increased importance with the evolution of resistance and may point to a potential role that CYP9A170 plays as a first line of defense against ingested insecticides.

The expression of digestive enzymes increased as larval development progressed. Similarly, amylases in *Helicoverpa armigera* [34] and a serine protease in *Spodoptera litura* [36] were expressed more as the larvae developed. Larvae likely require more energy and resources as they grow, corresponding to the rise in digestive enzyme expression to process food. In contrast, expression of detoxification related genes like CYP9A170 and mdr49 was relatively stable for most of larval development. Zhu et al. [38] reported that CYP9AQ1 and CYP9A3 expression in *Locusta migratoria* was consistent in the larval stages except for peaks in fifth and second instar respectively, which is reminiscent of CYP9A170 expression in *A. honmai*. Therefore, if a particular CYP is involved in insecticide resistance, it may be expressed throughout an individual’s life to prepare for insecticide exposure.

While the expression of different digestive enzymes has been shown to change according to diets [33, 34, 39, 40], few have explored gene expression patterns of digestive enzymes in response to food intake. Here, we found that *A. honmai* larvae did not regulate gene expression of several putative digestive enzymes within 4 hours of food intake after starvation. The feeding habit of lepidopteran larvae may explain the lack of response in gene expression to acute food intake in *A. honmai* larvae. In animals that feed continuously and *ad libitum*, there would be little need to retain a capacity for gene inducibility. It is important to emphasize that the quantitative relationship between mRNA, and proenzyme/ enzyme protein is complex, making it impossible to say if the changes in mRNA reflect intralumen enzyme activity. Depending on the enzyme, protein may be stored in vesicles prior to secretion, exported to the cell membrane, secreted into the lumen, and in some cases recovered via endocytosis for recycling [41]. Each step in the pathway from synthesis to lumen activity can be regulated independently.

Constitutive upregulation of detoxification enzymes has often been implicated in insecticide resistance, but induction of these enzymes upon insecticide exposure can provide the animal with additional protection [13, 42]. For example, permethrin-resistant mosquitoes *Culex quinquefasciatus* not only had higher constitutive levels of several CYPs, but they also induce additional CYP expression upon permethrin exposure [43]. On the other hand, there are also genes in resistant populations that are not constitutively upregulated but respond to toxin exposure, as is the case for multiple CYPs in permethrin resistance house flies *Musca domestica* [44]. In this study, *A. honmai* larvae had constitutively greater expression of CYP9A170, which is consistent with other tebufenozide resistant strains [12]. However, CYP9A170 expression did not respond to tebufenozide exposure. This result is unlikely to be due to ineffective tebufenozide treatment, as gene expression of the positive control, HR3, an indicator for tebufenozide exposure [23], responded in a dose-dependent manner in either strain. In addition, an ABC transporter gene, mdr49-like protein, was induced in a dose-dependent manner by tebufenozide interestingly only in the resistant strain. *mdr49* expression in *Drosophila melanogaster* responded to colchicine feeding [45], and several ABC transporter genes in *H. armigera* had distinct responses to different insecticides [46]. It is therefore probable that this particular ABC transporter gene is implicated in protecting *A. honmai* larvae during tebufenozide exposure. Future efforts charactering the function, basal levels, and responsiveness to insecticides of individual detoxification genes would greatly improve our understanding of the mechanisms of insecticide resistance.

Different strains of *A. honmai* resistant to tebufenozide share mechanisms that confer resistance; however, they also appear to each have distinct microevolutionary variations. Similar to other tebufenozide-resistant strains [12], our resistant larvae have the mutation on an ecdysone receptor that contributes to tebufenozide resistance [20], as well as increased expression of detoxification enzymes such as CYP9A170. Notably, our resistant strain also had higher expression of digestive enzymes than the susceptible strain. This observation was only present in select resistant *A. honmai* strain (Yui; Fig 6C). Therefore, populations that evolve resistance likely have specific microevolutionary differences beyond shared mechanisms that confer resistance. In support, only 55 genes out of 63, 454 were identified as common DEGs in a pairwise comparison between the two tebufenozide-susceptible and the two resistant *A. honmai* strains [12]. However, there can be up to 5194 DEGs in the comparison between the Met (R) and Kanaya (S) strains [12], and 1855 DEGs in this study, suggesting that resistant populations can differ from susceptible populations in aspects other than those that directly contribute to resistance.

The extent to which these microevolutionary variations in gene expression and physiology affect life-history remains unclear. However, other physiological systems have been implicated in the evolution of insecticide resistance and associated fitness costs. For example, pyrethroid-resistant *Sitophilus zeamais* with no fitness cost had greater body mass, respiratory rate, and digestive enzyme activity than a resistant counterpart with fitness cost [47, 48]. Similarly, malathion-resistant *Tribolium castaneum* that did not show fitness costs had structural and physiological differences in their digestive tract compared to their susceptible counterpart [49]. Therefore, digestive capabilities of the animal likely contribute to whether a population that evolves insecticide resistance also demonstrates fitness costs, especially when more resource intensive mechanisms of resistance, such as increased production of detoxification enzymes, are in question.

## Limitations of the study

We acknowledge caveats that limit generalization and interpretation of the results in this study. Most of this study relied on gene expression data to characterize genetic differences between our susceptible and resistant strains of *A. honmai*. The first caveat to consider is the biosynthetic pathway for proteins.

Although the strains differed at the transcript level, these differences may not culminate in changes in protein, either as precursor, total active enzyme, or intra-lumen catalytic activity. Ultimately, the important phenotype for any of the digestive genes is the capacity to digest the very specific molecules the enzyme targets. Furthermore, for any particular reaction, there are often several gene products that contribute to hydrolysis of a class of molecules, making it difficult to connect one step in the enzyme synthetic process (i.e., transcription) to a digestive phenotypic capacity. The follow-up question of most relevance to this study is whether a putative change in digestion contributes to a change in fitness and whether such changes can overcome evolutionary costs of maintaining resistance in the absence of pesticide, assuming the presence of a fitness cost. Nonetheless, differences in transcript profiles between evolved lines provide evidence of a genetic distinction between the lines.

Another important caveat to consider is the distinct evolutionary history of our two strains. The two strains have been isolated for close to seven decades. We do not know how each of the strains have evolved and adapted to their respective environments in the field and lab, only that they are different beyond the detoxification system, which has been traditionally associated with insecticide resistance. Future studies that compare digestive physiology between strains would address these issues by investigating differences at various levels of organization more comprehensively over the time course of digestion, while employing more closely related strains or even genetic manipulation to control for the genetic differences between strains to allow for better inference.

## Conclusion

Many differences can arise between strains when populations develop insecticide resistance. Here, we confirmed that our resistant *A. honmai* had greater expression of detoxification enzymes such as CYP9A170, which is similar to other *A. honmai* populations that are resistant to tebufenozide. We also demonstrated that methodological considerations for parameters such as tissues, development stage, and exposure to stimuli can affect the transcriptional patterns that are observed. For example, our resistant strain increased the expression of mdr49-like protein only when they were exposed to tebufenozide. At the same time, populations can have microevolutionary variations that are not directly related to insecticide resistance. In the case of our resistant population, larvae expressed more of multiple putative digestive genes, which may be associated with better digestive capabilities. Future studies on fitness costs of the evolution of insecticide resistance should consider and characterize other physiological traits in specific populations, in addition to direct mechanisms of resistance, to bridge the connection between physiology and life-history.

## Acknowledgements

Thank you to Mckenzie Macklin for help with data collection, and to Talia Fleming and Davide Bonifacio-Proietto for help with colony maintenance.

## Data Availability Statement

The raw RNA-seq reads have been deposited in NCBI’s Gene Expression Omnibus [50] and are accessible through GEO Series accession number GSE288080. Other experimental data have been deposited in Borealis (https://doi.org/10.5683/SP3/8JBFMN).

