## Supplementary information for "Exploring the connections between digestion and detoxification in microevolution of insecticide resistance of the tea tortrix moth, *Adoxophyes honmai*"

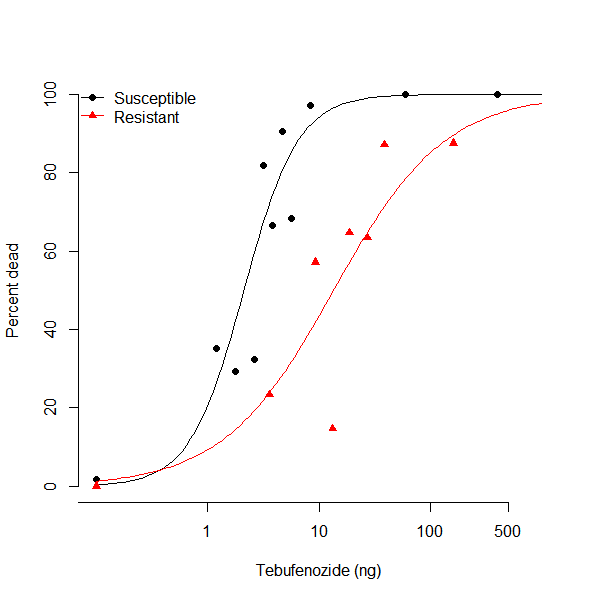

**Figure S1. Dose-dependent mortality (expressed in percent dead) of susceptible and resistant third instar larvae given a droplet of tebufenozide.**

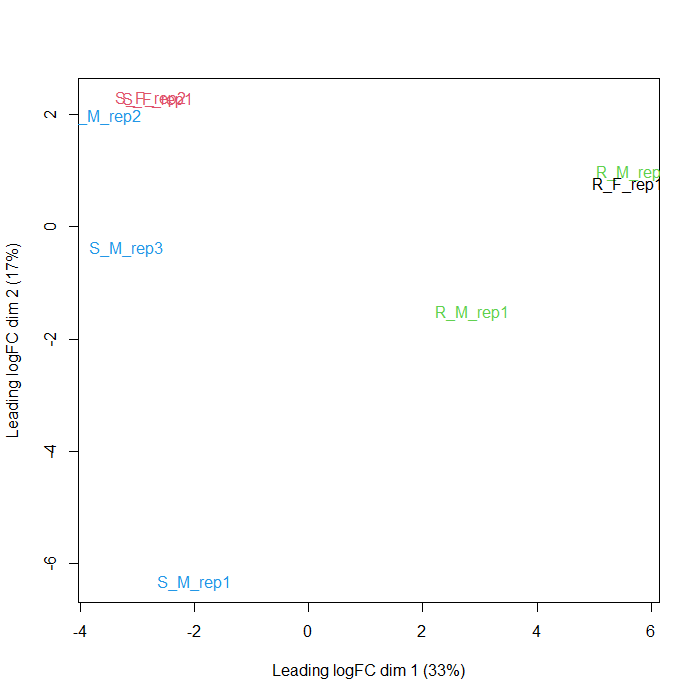

**Figure S2. Multidimentional scaling (MDS) plot visualization of the distance between the midgut transcriptomes of male (M) and female (F) susceptible (S) and resistant (R) strains of *A. honmai* (Fig 1). Males and females were grouped closer together than by strains.**

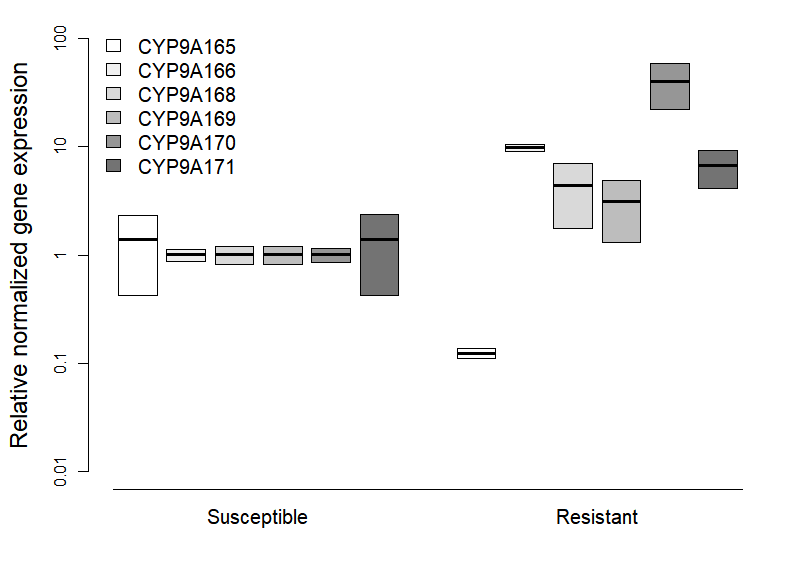

**Figure S3. Normalized relative fold-change in gene expression of cytochrome p450s associated with tebufenozide resistance (Uchibori-Asano et al. 2019) in third instar susceptible and resistant larvae. Expression values of each gene for each sample were normalized using the Cq of the reference gene, rp49, then divided by the mean normalized value of the respective gene for susceptible larvae to obtain the normalized relative fold-change.**

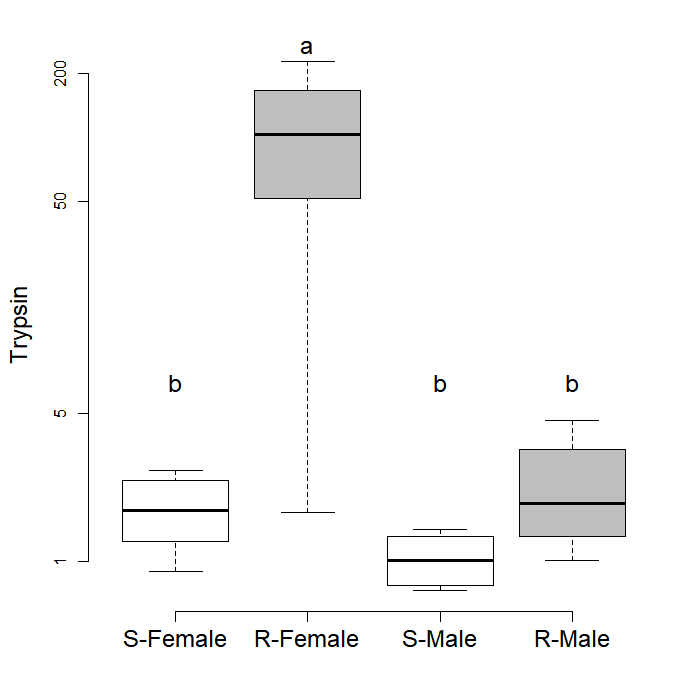

**Figure S4.** Normalized fold-change in trypsin expression in the digestive tract of fifth instar susceptible (S) and resistant (R) larvae relative to the mean in susceptible DT. For each sample, Cq values of each gene were first normalized using the Cq of the reference gene (rp49), then divided by the mean normalized value of the DT of susceptible larvae to obtain the normalized relative fold-change for each gene. S-Female: n = 5; R-Female: n = 4; S-Female: n = 4; R-Male: n = 4. Different letters indicate significant differences (p < 0.05) identified in the *post hoc* pairwise comparisons using Tukey’s test with Benjamini-Hochberg adjustment for multiple testing.

**Table S1. Primers for real time qPCR.**

| Target | Forward primer (5’-3’) | Reverse primer (5’-3’) |
| --- | --- | --- |
| rp49* | CCG TCA CAT GCT ACC CAA TG | CAG TAG GAC CTG TTC TGC ATC |
| CYP9A170* | GGC ACA GGA GAT CAG GGA AA | CTC GGA GAC AAC CAT GTC CA |
| Amylase | GAA AAC GTC GGT ACT GGA GG | CTC TAT CAG GAC AGC AGC CA |
| Maltase | TAC ATG TAC GGG AGA GCG TC | GCA GTT CTA CCT CCA CCA GT |
| Pancreatic lipase | AAC TTC AAC GCC AAT GTC CC | GCA AGC CTC CTC CAA TCA AG |
| Gastric lipase | TGG GTG TGG TTC CTT GAG AA | TCA TCT GCT GAC TTC GTG GT |
| Trypsin | CAT CTG TTA GCT GCG TCA CC | GGT CTG CTC TGC TGT ACT CA |
| Chymotrypsin | CGT GCA GAT CTC AAT GTG GG | AGA TCA CGT TGC CAG TCA TC |
| HR3 | CCT TGT GCT ATT GTG GCC AG | TCC AGA TGC ATG AGG GAC AG |
| mdr49 | AAT TCG GCG AAC TCA ACA CC | AAC TAG TCG CCT TCC TCC AC |

***Obtained from Uchibori-Asano et al. 2019.**

**Table S2. Linear models selection for the effects of strain and tissue on the expression of CYP9A170 in the digestive tract and the carcass of 5^th^ instar larvae (Fig 2B). The statistics for the model of best fit were bolded.**

| Model | ΔAICc | Weight |
| --- | --- | --- |
| **y~strain*tissue** | **0.00** | **1.00** |
| y~strain+tissue | 20.07 | 0.00 |
| y~strain | 19.42 | 0.00 |
| y~tissue | 73.21 | 0.00 |
| y~1 | 71.19 | 0.00 |

**Table S3. Post-hoc comparisons between strains and tissues for CYP9A170 expression in the best-fit model (Table S2).**

| Comparison | Estimate | Std. Error | t value | Pr(>\|t\|) |
| --- | --- | --- | --- | --- |
| R Carcass - R G == 0 | -1.1988 | 0.4559 | -2.629 | 0.0134* |
| S G - R G == 0 | -6.569 | 0.4431 | -14.826 | 1.47E-14*** |
| S Carcass - R G == 0 | -4.4101 | 0.4431 | -9.954 | 1.02E-10*** |
| S G - R Carcass == 0 | -5.3703 | 0.4431 | -12.121 | 1.30E-12*** |
| S Carcass - R Carcass == 0 | -3.2114 | 0.4431 | -7.248 | 6.82E-08*** |
| S Carcass - S G == 0 | 2.1589 | 0.4298 | 5.023 | 2.62E-05*** |

**Table S4. Detection limits, the Cq at which qPCR efficiency began to decrease based on a standard curve, and the mean Cq in the carcass for putative digestive enzymes.**

| Gene | Detection limit (Cq) | Mean carcass Cq |
| --- | --- | --- |
| Pancreatic lipase | 26.2 | 31.1 |
| Gastric lipase | 28.1 | 30.0 |
| Trypsin | 22.5 | 28.2 |
| Chymotrypsin | 29.5 | 33.0 |
| Amylase | 26.1 | 24.4 |
| Maltase | 27.3 | 31.2 |

**Table S5. Linear models selection for the effects of strain on the expression of putative digestive enzymes in the digestive tract of 5^th^ instar larvae (Fig 2C). The statistics for the model of best fit were bolded.**

| Model | Pancreatic lipase | | Gastric lipase | | Trypsin | | Amylase | | Maltase | | Chymotrypsin | |
| --- | --- | --- | --- | --- | --- | --- | --- | --- | --- | --- | --- | --- |
|  | ΔAICc | Weight | ΔAICc | Weight | ΔAICc | Weight | ΔAICc | Weight | ΔAICc | Weight | ΔAICc | Weight |
| y~strain*sex | 7.51 | 0.02 | 1.74 | 0.15 | **0.00** | **0.91** | 10.28 | 0.00 | 6.37 | 0.03 | 2.69 | 0.11 |
| y~strain | **0.00** | **0.98** | 1.44 | 0.17 | 5.54 | 0.06 | 2.99 | 0.18 | **0.00** | **0.75** | 2.03 | 0.15 |
| y~sex | 17.91 | 0.00 | 0.20 | 0.32 | 7.79 | 0.02 | 2.99 | 0.15 | 5.12 | 0.06 | 0.61 | 0.31 |
| y~1 | 14.92 | 0.00 | **0.00** | **0.67** | 8.79 | 0.01 | **0.00** | **0.69** | 2.99 | 0.18 | **0.00** | **0.42** |

**Table S6. Linear models selection for the effects of strain and feeding on the expression of putative digestive enzymes in 3^rd^ and 5^th^ instar larvae (Fig 3). The statistics for the model of best fit were bolded.**

| Model | Pancreatic lipase | | | | Trypsin | | | | Amylase | | | |
| --- | --- | --- | --- | --- | --- | --- | --- | --- | --- | --- | --- | --- |
|  | Third | | Fifth | | Third | | Fifth | | Third | | Fifth | |
|  | ΔAICc | Weight | ΔAICc | Weight | ΔAICc | Weight | ΔAICc | Weight | ΔAICc | Weight | ΔAICc | Weight |
| y~Strain*Feed | **0.00** | **0.68** | 3.49 | 0.10 | 3.70 | 0.11 | 3.77 | 0.09 | 3.26 | 0.08 | 5.21 | 0.05 |
| y~Strain+Feed | 3.33 | 0.13 | 0.93 | 0.35 | 2.34 | 0.21 | 1.17 | 0.33 | 2.41 | 0.12 | 2.54 | 0.18 |
| y~Feed | 4.14 | 0.09 | 25.54 | 0.00 | 37.58 | 0.00 | 28.25 | 0.00 | 2.94 | 0.09 | 5.89 | 0.03 |
| y~Strain | 4.87 | 0.06 | **0.00** | **0.55** | **0.00** | **0.68** | **0.00** | **0.59** | **0.00** | **0.41** | **0.00** | **0.63** |
| y~1 | 5.42 | 0.05 | 23.92 | 0.00 | 35.34 | 0.00 | 26.47 | 0.00 | 0.64 | 0.30 | 3.51 | 0.11 |

**Table S7.** **Post-hoc comparisons between fed and unfed larvae for pancreatic lipase expression in the best-fit model (Table S5).**

| Comparison | Estimate | Std. Error | t value | Pr(>\|t\|) |
| --- | --- | --- | --- | --- |
| R Fed - R Unfed == 0 | -0.05 | 0.64 | -0.07 | 0.94 |
| S Unfed - R Unfed == 0 | 0.22 | 0.69 | 0.32 | 0.90 |
| S Fed - R Unfed == 0 | -2.24 | 0.69 | -3.23 | <0.01** |
| S Unfed - R Fed == 0 | 0.27 | 0.69 | 0.39 | 0.90 |
| S Fed - R Fed == 0 | -2.20 | 0.69 | -3.17 | <0.01** |
| S Fed - S Unfed == 0 | -2.47 | 0.74 | -3.32 | <0.01** |

**Table S8. Linear models selection for the effects of strain and larval developmental stage on the expression of digestive and detoxification enzymes in larvae (Fig 4). The statistics for the model of best fit were bolded.**

| Model | Pancreatic lipase | | Amylase | | Trypsin | | Maltase | | CYP9A170 | | MDR | |
| --- | --- | --- | --- | --- | --- | --- | --- | --- | --- | --- | --- | --- |
|  | ΔAICc | Weight | ΔAICc | Weight | ΔAICc | Weight | ΔAICc | Weight | ΔAICc | Weight | ΔAICc | Weight |
| y~stage*strain | 17.19 | 0.00 | 13.92 | 0.00 | 3.92 | 0.12 | 3.94 | 0.12 | 14.73 | 0.00 | 18.52 | 0.00 |
| y~stage+strain | **0.00** | **1.00** | 2.30 | 0.24 | **0.00** | **0.86** | **0.00** | **0.88** | **0.00** | **1.00** | **0.00** | **1.00** |
| y~strain | 11.21 | 0.00 | 81.40 | 0.00 | 7.12 | 0.02 | 41.74 | 0.00 | 43.84 | 0.00 | 18.20 | 0.00 |
| y~stage | 32.51 | 0.00 | **0.00** | **0.76** | 188.24 | 0.00 | 42.64 | 0.00 | 213.69 | 0.00 | 17.92 | 0.00 |
| y~1 | 36.78 | 0.00 | 79.37 | 0.00 | 174.12 | 0.00 | 68.43 | 0.00 | 206.69 | 0.00 | 31.17 | 0.00 |

**Table S9.** **Post-hoc comparisons between developmental stages for pancreatic lipase expression in the best-fit model (Table S8).**

| Comparison | Estimate | Std. Error | t value | Pr(>\|t\|) |
| --- | --- | --- | --- | --- |
| 1.5 - 1 == 0 | -0.07333 | 1.01408 | -0.072 | 0.94248 |
| 2 - 1 == 0 | -0.73667 | 1.01408 | -0.726 | 0.6209 |
| 2.5 - 1 == 0 | -1.84 | 1.01408 | -1.814 | 0.17138 |
| 3 - 1 == 0 | -0.63083 | 1.01408 | -0.622 | 0.63442 |
| 3.5 - 1 == 0 | -1.05583 | 1.01408 | -1.041 | 0.46567 |
| 4 - 1 == 0 | 0.38583 | 1.01408 | 0.38 | 0.75464 |
| 4.5 - 1 == 0 | 1.24417 | 1.01408 | 1.227 | 0.40051 |
| 5 - 1 == 0 | 2.55667 | 1.01408 | 2.521 | 0.05377 |
| 5.5 - 1 == 0 | 1.92667 | 1.01408 | 1.9 | 0.15905 |
| 2 - 1.5 == 0 | -0.66333 | 1.01408 | -0.654 | 0.63442 |
| 2.5 - 1.5 == 0 | -1.76667 | 1.01408 | -1.742 | 0.18969 |
| 3 - 1.5 == 0 | -0.5575 | 1.01408 | -0.55 | 0.6734 |
| 3.5 - 1.5 == 0 | -0.9825 | 1.01408 | -0.969 | 0.48594 |
| 4 - 1.5 == 0 | 0.45917 | 1.01408 | 0.453 | 0.73305 |
| 4.5 - 1.5 == 0 | 1.3175 | 1.01408 | 1.299 | 0.37182 |
| 5 - 1.5 == 0 | 2.63 | 1.01408 | 2.593 | 0.05377 |
| 5.5 - 1.5 == 0 | 2 | 1.01408 | 1.972 | 0.15003 |
| 2.5 - 2 == 0 | -1.10333 | 1.01408 | -1.088 | 0.44838 |
| 3 - 2 == 0 | 0.10583 | 1.01408 | 0.104 | 0.93792 |
| 3.5 - 2 == 0 | -0.31917 | 1.01408 | -0.315 | 0.78861 |
| 4 - 2 == 0 | 1.1225 | 1.01408 | 1.107 | 0.44838 |
| 4.5 - 2 == 0 | 1.98083 | 1.01408 | 1.953 | 0.15003 |
| 5 - 2 == 0 | 3.29333 | 1.01408 | 3.248 | 0.01741 * |
| 5.5 - 2 == 0 | 2.66333 | 1.01408 | 2.626 | 0.05377 |
| 3 - 2.5 == 0 | 1.20917 | 1.01408 | 1.192 | 0.40795 |
| 3.5 - 2.5 == 0 | 0.78417 | 1.01408 | 0.773 | 0.60141 |
| 4 - 2.5 == 0 | 2.22583 | 1.01408 | 2.195 | 0.10485 |
| 4.5 - 2.5 == 0 | 3.08417 | 1.01408 | 3.041 | 0.02213 * |
| 5 - 2.5 == 0 | 4.39667 | 1.01408 | 4.336 | 0.00146 ** |
| 5.5 - 2.5 == 0 | 3.76667 | 1.01408 | 3.714 | 0.00727 ** |
| 3.5 - 3 == 0 | -0.425 | 1.01408 | -0.419 | 0.74192 |
| 4 - 3 == 0 | 1.01667 | 1.01408 | 1.003 | 0.47745 |
| 4.5 - 3 == 0 | 1.875 | 1.01408 | 1.849 | 0.16793 |
| 5 - 3 == 0 | 3.1875 | 1.01408 | 3.143 | 0.01937 * |
| 5.5 - 3 == 0 | 2.5575 | 1.01408 | 2.522 | 0.05377 |
| 4 - 3.5 == 0 | 1.44167 | 1.01408 | 1.422 | 0.32315 |
| 4.5 - 3.5 == 0 | 2.3 | 1.01408 | 2.268 | 0.09486 |
| 5 - 3.5 == 0 | 3.6125 | 1.01408 | 3.562 | 0.0082 ** |
| 5.5 - 3.5 == 0 | 2.9825 | 1.01408 | 2.941 | 0.02568 * |
| 4.5 - 4 == 0 | 0.85833 | 1.01408 | 0.846 | 0.56134 |
| 5 - 4 == 0 | 2.17083 | 1.01408 | 2.141 | 0.11098 |
| 5.5 - 4 == 0 | 1.54083 | 1.01408 | 1.519 | 0.28189 |
| 5 - 4.5 == 0 | 1.3125 | 1.01408 | 1.294 | 0.37182 |
| 5.5 - 4.5 == 0 | 0.6825 | 1.01408 | 0.673 | 0.63442 |
| 5.5 - 5 == 0 | -0.63 | 1.01408 | -0.621 | 0.63442 |

**Table S10.** **Post-hoc comparisons between developmental stages for trypsin expression in the best-fit model (Table S8).**

| Comparison | Estimate | Std. Error | t value | Pr(>\|t\|) |
| --- | --- | --- | --- | --- |
| 1.5 - 1 == 0 | 0.07333 | 0.62075 | 0.118 | 0.9268 |
| 2 - 1 == 0 | 0.7775 | 0.62075 | 1.253 | 0.3835 |
| 2.5 - 1 == 0 | 0.7475 | 0.62075 | 1.204 | 0.4 |
| 3 - 1 == 0 | 0.65 | 0.62075 | 1.047 | 0.446 |
| 3.5 - 1 == 0 | 1.11167 | 0.62075 | 1.791 | 0.214 |
| 4 - 1 == 0 | 1.80417 | 0.62075 | 2.906 | 0.0399* |
| 4.5 - 1 == 0 | 0.99667 | 0.62075 | 1.606 | 0.2384 |
| 5 - 1 == 0 | 2.0575 | 0.62075 | 3.315 | 0.0187* |
| 5.5 - 1 == 0 | 2.27833 | 0.62075 | 3.67 | 0.0127* |
| 2 - 1.5 == 0 | 0.70417 | 0.62075 | 1.134 | 0.4292 |
| 2.5 - 1.5 == 0 | 0.67417 | 0.62075 | 1.086 | 0.4342 |
| 3 - 1.5 == 0 | 0.57667 | 0.62075 | 0.929 | 0.5152 |
| 3.5 - 1.5 == 0 | 1.03833 | 0.62075 | 1.673 | 0.2273 |
| 4 - 1.5 == 0 | 1.73083 | 0.62075 | 2.788 | 0.0469* |
| 4.5 - 1.5 == 0 | 0.92333 | 0.62075 | 1.487 | 0.2735 |
| 5 - 1.5 == 0 | 1.98417 | 0.62075 | 3.196 | 0.0205* |
| 5.5 - 1.5 == 0 | 2.205 | 0.62075 | 3.552 | 0.0127* |
| 2.5 - 2 == 0 | -0.03 | 0.62075 | -0.048 | 0.9615 |
| 3 - 2 == 0 | -0.1275 | 0.62075 | -0.205 | 0.9143 |
| 3.5 - 2 == 0 | 0.33417 | 0.62075 | 0.538 | 0.7393 |
| 4 - 2 == 0 | 1.02667 | 0.62075 | 1.654 | 0.2273 |
| 4.5 - 2 == 0 | 0.21917 | 0.62075 | 0.353 | 0.8153 |
| 5 - 2 == 0 | 1.28 | 0.62075 | 2.062 | 0.1439 |
| 5.5 - 2 == 0 | 1.50083 | 0.62075 | 2.418 | 0.0864 |
| 3 - 2.5 == 0 | -0.0975 | 0.62075 | -0.157 | 0.9162 |
| 3.5 - 2.5 == 0 | 0.36417 | 0.62075 | 0.587 | 0.7393 |
| 4 - 2.5 == 0 | 1.05667 | 0.62075 | 1.702 | 0.2273 |
| 4.5 - 2.5 == 0 | 0.24917 | 0.62075 | 0.401 | 0.8153 |
| 5 - 2.5 == 0 | 1.31 | 0.62075 | 2.11 | 0.1439 |
| 5.5 - 2.5 == 0 | 1.53083 | 0.62075 | 2.466 | 0.0856 |
| 3.5 - 3 == 0 | 0.46167 | 0.62075 | 0.744 | 0.6254 |
| 4 - 3 == 0 | 1.15417 | 0.62075 | 1.859 | 0.197 |
| 4.5 - 3 == 0 | 0.34667 | 0.62075 | 0.558 | 0.7393 |
| 5 - 3 == 0 | 1.4075 | 0.62075 | 2.267 | 0.114 |
| 5.5 - 3 == 0 | 1.62833 | 0.62075 | 2.623 | 0.064 |
| 4 - 3.5 == 0 | 0.6925 | 0.62075 | 1.116 | 0.4292 |
| 4.5 - 3.5 == 0 | -0.115 | 0.62075 | -0.185 | 0.9143 |
| 5 - 3.5 == 0 | 0.94583 | 0.62075 | 1.524 | 0.2669 |
| 5.5 - 3.5 == 0 | 1.16667 | 0.62075 | 1.879 | 0.197 |
| 4.5 - 4 == 0 | -0.8075 | 0.62075 | -1.301 | 0.3676 |
| 5 - 4 == 0 | 0.25333 | 0.62075 | 0.408 | 0.8153 |
| 5.5 - 4 == 0 | 0.47417 | 0.62075 | 0.764 | 0.6254 |
| 5 - 4.5 == 0 | 1.06083 | 0.62075 | 1.709 | 0.2273 |
| 5.5 - 4.5 == 0 | 1.28167 | 0.62075 | 2.065 | 0.1439 |
| 5.5 - 5 == 0 | 0.22083 | 0.62075 | 0.356 | 0.8153 |

**Table S11.** **Post-hoc comparisons between developmental stages for amylase expression in the best-fit model (Table S8).**

| Comparison | Estimate | Std. Error | t value | Pr(>\|t\|) |
| --- | --- | --- | --- | --- |
| 1.5 - 1 == 0 | 0.05 | 0.4368 | 0.114 | 0.929737 |
| 2 - 1 == 0 | -0.8592 | 0.4368 | -1.967 | 0.086182 |
| 2.5 - 1 == 0 | -0.705 | 0.4368 | -1.614 | 0.164091 |
| 3 - 1 == 0 | -0.8767 | 0.4368 | -2.007 | 0.081697 |
| 3.5 - 1 == 0 | -0.2475 | 0.4368 | -0.567 | 0.643647 |
| 4 - 1 == 0 | 0.3358 | 0.4368 | 0.769 | 0.539557 |
| 4.5 - 1 == 0 | 0.9967 | 0.4368 | 2.282 | 0.052351 |
| 5 - 1 == 0 | 0.8875 | 0.4368 | 2.032 | 0.080251 |
| 5.5 - 1 == 0 | 3.2283 | 0.4368 | 7.391 | 2.73e-10 *** |
| 2 - 1.5 == 0 | -0.9092 | 0.4368 | -2.081 | 0.074471 |
| 2.5 - 1.5 == 0 | -0.755 | 0.4368 | -1.728 | 0.134553 |
| 3 - 1.5 == 0 | -0.9267 | 0.4368 | -2.121 | 0.070686 |
| 3.5 - 1.5 == 0 | -0.2975 | 0.4368 | -0.681 | 0.58885 |
| 4 - 1.5 == 0 | 0.2858 | 0.4368 | 0.654 | 0.593351 |
| 4.5 - 1.5 == 0 | 0.9467 | 0.4368 | 2.167 | 0.066216 |
| 5 - 1.5 == 0 | 0.8375 | 0.4368 | 1.917 | 0.092876 |
| 5.5 - 1.5 == 0 | 3.1783 | 0.4368 | 7.276 | 4.04e-10 *** |
| 2.5 - 2 == 0 | 0.1542 | 0.4368 | 0.353 | 0.77658 |
| 3 - 2 == 0 | -0.0175 | 0.4368 | -0.04 | 0.968115 |
| 3.5 - 2 == 0 | 0.6117 | 0.4368 | 1.4 | 0.223953 |
| 4 - 2 == 0 | 1.195 | 0.4368 | 2.736 | 0.018143 * |
| 4.5 - 2 == 0 | 1.8558 | 0.4368 | 4.249 | 0.000185 *** |
| 5 - 2 == 0 | 1.7467 | 0.4368 | 3.999 | 0.000400 *** |
| 5.5 - 2 == 0 | 4.0875 | 0.4368 | 9.358 | 2.50e-14 *** |
| 3 - 2.5 == 0 | -0.1717 | 0.4368 | -0.393 | 0.762889 |
| 3.5 - 2.5 == 0 | 0.4575 | 0.4368 | 1.047 | 0.371526 |
| 4 - 2.5 == 0 | 1.0408 | 0.4368 | 2.383 | 0.042514 * |
| 4.5 - 2.5 == 0 | 1.7017 | 0.4368 | 3.896 | 0.000542 *** |
| 5 - 2.5 == 0 | 1.5925 | 0.4368 | 3.646 | 0.001226 ** |
| 5.5 - 2.5 == 0 | 3.9333 | 0.4368 | 9.005 | 1.13e-13 *** |
| 3.5 - 3 == 0 | 0.6292 | 0.4368 | 1.44 | 0.214593 |
| 4 - 3 == 0 | 1.2125 | 0.4368 | 2.776 | 0.017133 * |
| 4.5 - 3 == 0 | 1.8733 | 0.4368 | 4.289 | 0.000174 *** |
| 5 - 3 == 0 | 1.7642 | 0.4368 | 4.039 | 0.000374 *** |
| 5.5 - 3 == 0 | 4.105 | 0.4368 | 9.398 | 2.50e-14 *** |
| 4 - 3.5 == 0 | 0.5833 | 0.4368 | 1.335 | 0.244171 |
| 4.5 - 3.5 == 0 | 1.2442 | 0.4368 | 2.848 | 0.014756 * |
| 5 - 3.5 == 0 | 1.135 | 0.4368 | 2.598 | 0.025220 * |
| 5.5 - 3.5 == 0 | 3.4758 | 0.4368 | 7.957 | 1.92e-11 *** |
| 4.5 - 4 == 0 | 0.6608 | 0.4368 | 1.513 | 0.193323 |
| 5 - 4 == 0 | 0.5517 | 0.4368 | 1.263 | 0.269067 |
| 5.5 - 4 == 0 | 2.8925 | 0.4368 | 6.622 | 8.67e-09 *** |
| 5 - 4.5 == 0 | -0.1092 | 0.4368 | -0.25 | 0.840469 |
| 5.5 - 4.5 == 0 | 2.2317 | 0.4368 | 5.109 | 6.87e-06 *** |
| 5.5 - 5 == 0 | 2.3408 | 0.4368 | 5.359 | 2.62e-06 *** |

**Table S12.** **Post-hoc comparisons between developmental stages for maltase expression in the best-fit model (Table S8).**

| Comparison | Estimate | Std. Error | t value | Pr(>\|t\|) |
| --- | --- | --- | --- | --- |
| 1.5 - 1 == 0 | -0.97667 | 0.44914 | -2.175 | 0.064829 |
| 2 - 1 == 0 | -1.615 | 0.44914 | -3.596 | 0.002193 ** |
| 2.5 - 1 == 0 | -1.85833 | 0.44914 | -4.138 | 0.000446 *** |
| 3 - 1 == 0 | -1.79083 | 0.44914 | -3.987 | 0.000682 *** |
| 3.5 - 1 == 0 | -1.62917 | 0.44914 | -3.627 | 0.002186 ** |
| 4 - 1 == 0 | -1.39667 | 0.44914 | -3.11 | 0.008274 ** |
| 4.5 - 1 == 0 | -0.62083 | 0.44914 | -1.382 | 0.254566 |
| 5 - 1 == 0 | -0.53 | 0.44914 | -1.18 | 0.349188 |
| 5.5 - 1 == 0 | 0.96917 | 0.44914 | 2.158 | 0.064829 |
| 2 - 1.5 == 0 | -0.63833 | 0.44914 | -1.421 | 0.245327 |
| 2.5 - 1.5 == 0 | -0.88167 | 0.44914 | -1.963 | 0.097857 |
| 3 - 1.5 == 0 | -0.81417 | 0.44914 | -1.813 | 0.125696 |
| 3.5 - 1.5 == 0 | -0.6525 | 0.44914 | -1.453 | 0.239713 |
| 4 - 1.5 == 0 | -0.42 | 0.44914 | -0.935 | 0.465603 |
| 4.5 - 1.5 == 0 | 0.35583 | 0.44914 | 0.792 | 0.537411 |
| 5 - 1.5 == 0 | 0.44667 | 0.44914 | 0.995 | 0.439338 |
| 5.5 - 1.5 == 0 | 1.94583 | 0.44914 | 4.332 | 0.000247 *** |
| 2.5 - 2 == 0 | -0.24333 | 0.44914 | -0.542 | 0.704903 |
| 3 - 2 == 0 | -0.17583 | 0.44914 | -0.391 | 0.764119 |
| 3.5 - 2 == 0 | -0.01417 | 0.44914 | -0.032 | 0.974895 |
| 4 - 2 == 0 | 0.21833 | 0.44914 | 0.486 | 0.706341 |
| 4.5 - 2 == 0 | 0.99417 | 0.44914 | 2.214 | 0.062026 |
| 5 - 2 == 0 | 1.085 | 0.44914 | 2.416 | 0.041128 * |
| 5.5 - 2 == 0 | 2.58417 | 0.44914 | 5.754 | 9.13e-07 *** |
| 3 - 2.5 == 0 | 0.0675 | 0.44914 | 0.15 | 0.900833 |
| 3.5 - 2.5 == 0 | 0.22917 | 0.44914 | 0.51 | 0.704903 |
| 4 - 2.5 == 0 | 0.46167 | 0.44914 | 1.028 | 0.430696 |
| 4.5 - 2.5 == 0 | 1.2375 | 0.44914 | 2.755 | 0.019334 * |
| 5 - 2.5 == 0 | 1.32833 | 0.44914 | 2.958 | 0.012224 * |
| 5.5 - 2.5 == 0 | 2.8275 | 0.44914 | 6.295 | 2.96e-07 *** |
| 3.5 - 3 == 0 | 0.16167 | 0.44914 | 0.36 | 0.770979 |
| 4 - 3 == 0 | 0.39417 | 0.44914 | 0.878 | 0.491252 |
| 4.5 - 3 == 0 | 1.17 | 0.44914 | 2.605 | 0.027714 * |
| 5 - 3 == 0 | 1.26083 | 0.44914 | 2.807 | 0.017763 * |
| 5.5 - 3 == 0 | 2.76 | 0.44914 | 6.145 | 3.00e-07 *** |
| 4 - 3.5 == 0 | 0.2325 | 0.44914 | 0.518 | 0.704903 |
| 4.5 - 3.5 == 0 | 1.00833 | 0.44914 | 2.245 | 0.060262 |
| 5 - 3.5 == 0 | 1.09917 | 0.44914 | 2.447 | 0.039976 * |
| 5.5 - 3.5 == 0 | 2.59833 | 0.44914 | 5.785 | 9.13e-07 *** |
| 4.5 - 4 == 0 | 0.77583 | 0.44914 | 1.727 | 0.144883 |
| 5 - 4 == 0 | 0.86667 | 0.44914 | 1.93 | 0.101253 |
| 5.5 - 4 == 0 | 2.36583 | 0.44914 | 5.268 | 6.32e-06 *** |
| 5 - 4.5 == 0 | 0.09083 | 0.44914 | 0.202 | 0.879181 |
| 5.5 - 4.5 == 0 | 1.59 | 0.44914 | 3.54 | 0.002411 ** |
| 5.5 - 5 == 0 | 1.49917 | 0.44914 | 3.338 | 0.004334 ** |

**Table S13.** **Post-hoc comparisons between developmental stages for CYP9A170 expression in the best-fit model (Table S8).**

| Comparison | Estimate | Std. Error | t value | Pr(>\|t\|) |
| --- | --- | --- | --- | --- |
| 1.5 - 1 == 0 | -1.8817 | 0.4462 | -4.217 | 0.000289 *** |
| 2 - 1 == 0 | -2.0142 | 0.4462 | -4.514 | 0.000104 *** |
| 2.5 - 1 == 0 | -2.3617 | 0.4462 | -5.292 | 5.68e-06 *** |
| 3 - 1 == 0 | -2.6017 | 0.4462 | -5.83 | 8.81e-07 *** |
| 3.5 - 1 == 0 | -2.2583 | 0.4462 | -5.061 | 1.28e-05 *** |
| 4 - 1 == 0 | -3.2092 | 0.4462 | -7.192 | 3.83e-09 *** |
| 4.5 - 1 == 0 | -2.5992 | 0.4462 | -5.825 | 8.81e-07 *** |
| 5 - 1 == 0 | -1.8667 | 0.4462 | -4.183 | 0.000292 *** |
| 5.5 - 1 == 0 | -0.8458 | 0.4462 | -1.895 | 0.124116 |
| 2 - 1.5 == 0 | -0.1325 | 0.4462 | -0.297 | 0.82188 |
| 2.5 - 1.5 == 0 | -0.48 | 0.4462 | -1.076 | 0.400017 |
| 3 - 1.5 == 0 | -0.72 | 0.4462 | -1.613 | 0.19169 |
| 3.5 - 1.5 == 0 | -0.3767 | 0.4462 | -0.844 | 0.53003 |
| 4 - 1.5 == 0 | -1.3275 | 0.4462 | -2.975 | 0.010830 * |
| 4.5 - 1.5 == 0 | -0.7175 | 0.4462 | -1.608 | 0.19169 |
| 5 - 1.5 == 0 | 0.015 | 0.4462 | 0.034 | 0.995365 |
| 5.5 - 1.5 == 0 | 1.0358 | 0.4462 | 2.321 | 0.055333 |
| 2.5 - 2 == 0 | -0.3475 | 0.4462 | -0.779 | 0.543214 |
| 3 - 2 == 0 | -0.5875 | 0.4462 | -1.317 | 0.288942 |
| 3.5 - 2 == 0 | -0.2442 | 0.4462 | -0.547 | 0.67011 |
| 4 - 2 == 0 | -1.195 | 0.4462 | -2.678 | 0.024058 * |
| 4.5 - 2 == 0 | -0.585 | 0.4462 | -1.311 | 0.288942 |
| 5 - 2 == 0 | 0.1475 | 0.4462 | 0.331 | 0.81398 |
| 5.5 - 2 == 0 | 1.1683 | 0.4462 | 2.618 | 0.026728 * |
| 3 - 2.5 == 0 | -0.24 | 0.4462 | -0.538 | 0.67011 |
| 3.5 - 2.5 == 0 | 0.1033 | 0.4462 | 0.232 | 0.855325 |
| 4 - 2.5 == 0 | -0.8475 | 0.4462 | -1.899 | 0.124116 |
| 4.5 - 2.5 == 0 | -0.2375 | 0.4462 | -0.532 | 0.67011 |
| 5 - 2.5 == 0 | 0.495 | 0.4462 | 1.109 | 0.39158 |
| 5.5 - 2.5 == 0 | 1.5158 | 0.4462 | 3.397 | 0.003571 ** |
| 3.5 - 3 == 0 | 0.3433 | 0.4462 | 0.769 | 0.543214 |
| 4 - 3 == 0 | -0.6075 | 0.4462 | -1.361 | 0.283182 |
| 4.5 - 3 == 0 | 0.0025 | 0.4462 | 0.006 | 0.99554 |
| 5 - 3 == 0 | 0.735 | 0.4462 | 1.647 | 0.19169 |
| 5.5 - 3 == 0 | 1.7558 | 0.4462 | 3.935 | 0.000613 *** |
| 4 - 3.5 == 0 | -0.9508 | 0.4462 | -2.131 | 0.079552 |
| 4.5 - 3.5 == 0 | -0.3408 | 0.4462 | -0.764 | 0.543214 |
| 5 - 3.5 == 0 | 0.3917 | 0.4462 | 0.878 | 0.520956 |
| 5.5 - 3.5 == 0 | 1.4125 | 0.4462 | 3.165 | 0.006952 ** |
| 4.5 - 4 == 0 | 0.61 | 0.4462 | 1.367 | 0.283182 |
| 5 - 4 == 0 | 1.3425 | 0.4462 | 3.008 | 0.010483 * |
| 5.5 - 4 == 0 | 2.3633 | 0.4462 | 5.296 | 5.68e-06 *** |
| 5 - 4.5 == 0 | 0.7325 | 0.4462 | 1.641 | 0.19169 |
| 5.5 - 4.5 == 0 | 1.7533 | 0.4462 | 3.929 | 0.000613 *** |
| 5.5 - 5 == 0 | 1.0208 | 0.4462 | 2.288 | 0.057052 |

**Table S14.** **Post-hoc comparisons between developmental stages for MDR expression in the best-fit model (Table S8).**

| Comparison | Estimate | Std. Error | t value | Pr(>\|t\|) |
| --- | --- | --- | --- | --- |
| 1.5 - 1 == 0 | 1.7875 | 0.5313 | 3.364 | 0.006813 ** |
| 2 - 1 == 0 | 2.43167 | 0.5313 | 4.577 | 0.000141 *** |
| 2.5 - 1 == 0 | 1.12917 | 0.5313 | 2.125 | 0.094824 |
| 3 - 1 == 0 | 2.50083 | 0.5313 | 4.707 | 0.000123 *** |
| 3.5 - 1 == 0 | 2.4875 | 0.5313 | 4.682 | 0.000123 *** |
| 4 - 1 == 0 | 2.54083 | 0.5313 | 4.782 | 0.000123 *** |
| 4.5 - 1 == 0 | 1.93917 | 0.5313 | 3.65 | 0.003641 ** |
| 5 - 1 == 0 | 1.79833 | 0.5313 | 3.385 | 0.006813 ** |
| 5.5 - 1 == 0 | 1.27917 | 0.5313 | 2.408 | 0.066514 |
| 2 - 1.5 == 0 | 0.64417 | 0.5313 | 1.212 | 0.365952 |
| 2.5 - 1.5 == 0 | -0.65833 | 0.5313 | -1.239 | 0.363289 |
| 3 - 1.5 == 0 | 0.71333 | 0.5313 | 1.343 | 0.363289 |
| 3.5 - 1.5 == 0 | 0.7 | 0.5313 | 1.318 | 0.363289 |
| 4 - 1.5 == 0 | 0.75333 | 0.5313 | 1.418 | 0.363289 |
| 4.5 - 1.5 == 0 | 0.15167 | 0.5313 | 0.285 | 0.937249 |
| 5 - 1.5 == 0 | 0.01083 | 0.5313 | 0.02 | 0.983769 |
| 5.5 - 1.5 == 0 | -0.50833 | 0.5313 | -0.957 | 0.451062 |
| 2.5 - 2 == 0 | -1.3025 | 0.5313 | -2.452 | 0.064693 |
| 3 - 2 == 0 | 0.06917 | 0.5313 | 0.13 | 0.983769 |
| 3.5 - 2 == 0 | 0.05583 | 0.5313 | 0.105 | 0.983769 |
| 4 - 2 == 0 | 0.10917 | 0.5313 | 0.205 | 0.966448 |
| 4.5 - 2 == 0 | -0.4925 | 0.5313 | -0.927 | 0.457704 |
| 5 - 2 == 0 | -0.63333 | 0.5313 | -1.192 | 0.365952 |
| 5.5 - 2 == 0 | -1.1525 | 0.5313 | -2.169 | 0.090676 |
| 3 - 2.5 == 0 | 1.37167 | 0.5313 | 2.582 | 0.053758 |
| 3.5 - 2.5 == 0 | 1.35833 | 0.5313 | 2.557 | 0.053758 |
| 4 - 2.5 == 0 | 1.41167 | 0.5313 | 2.657 | 0.051011 |
| 4.5 - 2.5 == 0 | 0.81 | 0.5313 | 1.525 | 0.325667 |
| 5 - 2.5 == 0 | 0.66917 | 0.5313 | 1.259 | 0.363289 |
| 5.5 - 2.5 == 0 | 0.15 | 0.5313 | 0.282 | 0.937249 |
| 3.5 - 3 == 0 | -0.01333 | 0.5313 | -0.025 | 0.983769 |
| 4 - 3 == 0 | 0.04 | 0.5313 | 0.075 | 0.983769 |
| 4.5 - 3 == 0 | -0.56167 | 0.5313 | -1.057 | 0.425005 |
| 5 - 3 == 0 | -0.7025 | 0.5313 | -1.322 | 0.363289 |
| 5.5 - 3 == 0 | -1.22167 | 0.5313 | -2.299 | 0.074721 |
| 4 - 3.5 == 0 | 0.05333 | 0.5313 | 0.1 | 0.983769 |
| 4.5 - 3.5 == 0 | -0.54833 | 0.5313 | -1.032 | 0.427966 |
| 5 - 3.5 == 0 | -0.68917 | 0.5313 | -1.297 | 0.363289 |
| 5.5 - 3.5 == 0 | -1.20833 | 0.5313 | -2.274 | 0.074721 |
| 4.5 - 4 == 0 | -0.60167 | 0.5313 | -1.132 | 0.389905 |
| 5 - 4 == 0 | -0.7425 | 0.5313 | -1.398 | 0.363289 |
| 5.5 - 4 == 0 | -1.26167 | 0.5313 | -2.375 | 0.066853 |
| 5 - 4.5 == 0 | -0.14083 | 0.5313 | -0.265 | 0.937249 |
| 5.5 - 4.5 == 0 | -0.66 | 0.5313 | -1.242 | 0.363289 |
| 5.5 - 5 == 0 | -0.51917 | 0.5313 | -0.977 | 0.450894 |

**Table S15.** **Linear models selection for the effects of strain, time, and tebufenozide dosage on the expression of HR3 and detoxification enzymes in larvae (Fig 5). The statistics for the model of best fit were bolded.**

| Model | HR3 | | CYP9A170 | | mdr49 | |
| --- | --- | --- | --- | --- | --- | --- |
|  | ΔAICc | Weight | ΔAICc | Weight | ΔAICc | Weight |
| y~strain*dose*time | 24.53 | 0.00 | 37.86 | 0.00 | 28.68 | 0.00 |
| y~strain*dose+time | 0.19 | 0.44 | 9.00 | 0.01 | 4.61 | 0.06 |
| y~strain+dose*time | 14.63 | 0.00 | 15.87 | 0.00 | 13.25 | 0.00 |
| y~strain*time+dose | 11.98 | 0.00 | 11.25 | 0.00 | 10.46 | 0.00 |
| y~strain+dose+time | 7.35 | 0.01 | 5.84 | 0.03 | 8.03 | 0.01 |
| y~strain*dose | **0.00** | **0.48** | 6.88 | 0.02 | **0.00** | **0.55** |
| y~strain+dose | 6.52 | 0.02 | 4.04 | 0.08 | 3.96 | 0.08 |
| y~dose*time | 32.05 | 0.00 | 89.24 | 0.00 | 24.93 | 0.00 |
| y~dose+time | 23.88 | 0.00 | 78.36 | 0.00 | 19.62 | 0.00 |
| y~strain*time | 49.49 | 0.00 | 6.38 | 0.02 | 7.17 | 0.02 |
| y~strain+time | 44.77 | 0.00 | 1.43 | 0.28 | 5.51 | 0.04 |
| y~strain | 42.03 | 0.00 | **0.00** | **0.57** | 1.76 | 0.23 |
| y~time | 51.90 | 0.00 | 73.58 | 0.00 | 16.74 | 0.00 |
| y~dose | 22.00 | 0.00 | 74.25 | 0.00 | 15.53 | 0.00 |
| y~strain+dose+time+strain:dose+strain:time | 5.08 | 0.04 | 14.94 | 0.00 | 6.73 | 0.02 |
| y~strain+dose+time+strain:dose+dose:time | 7.34 | 0.01 | 20.10 | 0.00 | 11.09 | 0.00 |
| y~strain+dose+time+strain:time+dose:time | 20.33 | 0.00 | 22.43 | 0.00 | 16.58 | 0.00 |
| y~1 | 49.02 | 0.00 | 69.86 | 0.00 | 12.99 | 0.00 |

**Table S16. Post-hoc comparisons between strains and tebufenozide dosage for HR3 expression in the best-fit model (Table S8).**

| Comparison | Estimate | Std. Error | t value | Pr(>\|t\|) |
| --- | --- | --- | --- | --- |
| R_1.2 ng - R_0 ng == 0 | 1.28 | 0.84 | 1.52 | 0.16 |
| R_8 ng - R_0 ng == 0 | 2.87 | 0.84 | 3.4 | 0.01 ** |
| S_0 ng - R_0 ng == 0 | 0.1 | 0.84 | 0.12 | 0.9 |
| S_1.2 ng - R_0 ng == 0 | 4.21 | 0.84 | 4.98 | <0.001 *** |
| S_8 ng - R_0 ng == 0 | 6.92 | 0.84 | 8.2 | <0.001 *** |
| R_8 ng - R_1.2 ng == 0 | 1.58 | 0.84 | 1.88 | 0.09 |
| S_0 ng - R_1.2 ng == 0 | -1.18 | 0.84 | -1.4 | 0.18 |
| S_1.2 ng - R_1.2 ng == 0 | 2.92 | 0.84 | 3.46 | <0.01 ** |
| S_8 ng - R_1.2 ng == 0 | 5.64 | 0.84 | 6.68 | <0.001 *** |
| S_0 ng - R_8 ng == 0 | -2.76 | 0.84 | -3.27 | <0.01 ** |
| S_1.2 ng - R_8 ng == 0 | 1.34 | 0.84 | 1.59 | 0.15 |
| S_8 ng - R_8 ng == 0 | 4.06 | 0.84 | 4.8 | <0.001 *** |
| S_1.2 ng - S_0 ng == 0 | 4.1 | 0.84 | 4.86 | <0.001 *** |
| S_8 ng - S_0 ng == 0 | 6.82 | 0.84 | 8.08 | <0.001 *** |
| S_8 ng - S_1.2 ng == 0 | 2.72 | 0.84 | 3.22 | <0.01 ** |

**Table S17.** **Post-hoc comparisons between strains and tebufenozide dosage for mdr49 expression in the best-fit model (Table S8).**

| Comparison | Estimate | Std. Error | t value | Pr(>\|t\|) |
| --- | --- | --- | --- | --- |
| R_1.2 ng - R_0 ng == 0 | 1.88 | 0.57 | 3.27 | <0.01 ** |
| R_8 ng - R_0 ng == 0 | 0.77 | 0.57 | 1.34 | 0.29 |
| S_0 ng - R_0 ng == 0 | -0.21 | 0.56 | -0.38 | 0.76 |
| S_1.2 ng - R_0 ng == 0 | -0.76 | 0.57 | -1.32 | 0.29 |
| S_8 ng - R_0 ng == 0 | -0.63 | 0.56 | -1.13 | 0.36 |
| R_8 ng - R_1.2 ng == 0 | -1.11 | 0.59 | -1.87 | 0.14 |
| S_0 ng - R_1.2 ng == 0 | -2.09 | 0.57 | -3.64 | <0.001 *** |
| S_1.2 ng - R_1.2 ng == 0 | -2.63 | 0.59 | -4.46 | <0.001 *** |
| S_8 ng - R_1.2 ng == 0 | -2.51 | 0.57 | -4.37 | <0.001 *** |
| S_0 ng - R_8 ng == 0 | -0.98 | 0.57 | -1.71 | 0.18 |
| S_1.2 ng - R_8 ng == 0 | -1.53 | 0.59 | -2.59 | 0.04 * |
| S_8 ng - R_8 ng == 0 | -1.4 | 0.57 | -2.44 | 0.046 * |
| S_1.2 ng - S_0 ng == 0 | -0.55 | 0.57 | -0.95 | 0.43 |
| S_8 ng - S_0 ng == 0 | -0.42 | 0.56 | -0.76 | 0.52 |
| S_8 ng - S_1.2 ng == 0 | 0.13 | 0.57 | 0.22 | 0.83 |
